# Single cell mutational profiling delineates clonal trajectories in myeloid malignancies

**DOI:** 10.1101/2020.02.07.938860

**Authors:** Linde A. Miles, Robert L. Bowman, Tiffany R. Merlinsky, Isabelle S. Csete, Aik Ooi, Robert Durruthy-Durruthy, Michael Bowman, Christopher Famulare, Minal A. Patel, Pedro Mendez, Chrysanthi Ainali, Mani Manivannan, Sombeet Sahu, Aaron D. Goldberg, Kelly Bolton, Ahmet Zehir, Raajit Rampal, Martin P. Carroll, Sara E. Meyer, Aaron D. Viny, Ross L. Levine

## Abstract

Myeloid malignancies, including acute myeloid leukemia (AML), arise from the proliferation and expansion of hematopoietic stem and progenitor cells which acquire somatic mutations. Bulk molecular profiling studies on patient samples have suggested that somatic mutations are obtained in a step-wise fashion, where mutant genes with high variant allele frequencies (VAFs) are proposed to occur early in disease development and mutations with lower VAFs are thought to be acquired later in disease progression^1–3^. Although bulk sequencing informs leukemia biology and prognostication, it cannot distinguish which mutations occur in the same clone(s), accurately measure clonal complexity and clone size, or offer definitive evidence of mutational order. To elucidate the clonal framework of myeloid malignancies, we performed single cell mutational profiling on 146 samples from 123 patients. We found AML is most commonly comprised of a small number of dominant clones, which in many cases harbor co-occurring mutations in epigenetic regulators. Conversely, mutations in signaling genes often occur more than once in distinct subclones consistent with increasing clonal diversity. We also used these data to map the clonal trajectory of each patient and found that specific mutation combinations (*FLT3-ITD* + *NPM1*^c^) synergize to promote clonal expansion and dominance. We combined cell surface protein expression with single cell mutational analysis to map somatic genotype and clonal architecture with immunophenotype. Our studies of clonal architecture at a single cell level provide novel insights into the pathogenesis of myeloid transformation and how clonal complexity contributes to disease progression.

## Results

The genomic landscape of myeloid malignancies has been well described, with a near complete catalogue of putative mutant driver genes^3–7^. While combinations of mutations have been functionally investigated in mouse models of disease, there remains uncertainty about the co-occurrence of somatic mutations within the same clone. The composition and combinatorial distribution of mutations across cell types and states continues to be poorly understood. To analyze the clonal architecture in clinical isolates from patients with myeloid malignancies, we performed single cell DNA sequencing on the Tapestri platform using a custom 109 amplicon panel covering 31 of the most frequently mutated genes in myeloid malignancies (Extended Table 1)^8^. We sequenced 740,529 cells from 146 samples from 123 patients with myeloid malignancies, including clonal hematopoiesis (CH), myeloproliferative neoplasms (MPN), and AML (Figure 1A). Samples from patients at diagnosis and at relapse were queried, with the majority of samples being from patients with relapsed/refractory disease (Figure 1B; Extended Table 2). The most common mutations identified in single cell profiling were found in *DNMT3A* (n=62 patients), *TET2* (n=58 patients), *NPM1* (n=37 patients) and *FLT3* (n=32 patients) genes, consistent with previous bulk sequencing studies^3–5^ (Extended Figure 1A-B). Mutational analysis identified that 80% of patients had at least 3 mutations in or near protein coding regions (Extended Figure 1C-D). We correlated single cell mutational findings with bulk sequencing results and found significant concordance between bulk sequencing and single cell sequencing results (Pearson correlation coefficient = 0.83; p ≤ 2.2 × 10^−16^; Extended Figure 1E). We next investigated mutational and clonal diversity in different disease subtypes, subdividing the AML into AML subclasses with epigenetic regulator mutations (DTAI; *DNMT3A, TET2, ASXL1* and/or *IDH1/2*) and DTAI-AML with signaling effector mutations, *RAS* and/or *FLT3*. The number of mutations per sample increased significantly from CH to MPN and then AML (Figure 1C; *FDR p* ≤ 7.15−10^− 6^ - 0.067 for all indicated comparisons). We observed a further increase in the number of mutations in AML cases with signaling effector mutations, specifically *RAS* and *FLT3* (*FDR p* ≤ 0.075). We next explored mutational diversity and clonal repertoire. Cells were grouped into clones if they possessed identical protein encoding SNVs, and a bootstrapping approach was applied to identify clones that were observed in at least 10 cells (Extended Figure 2). We were then able to enumerate the clones present in each sample (Figure 1D), and at what frequency they were observed. We then used this data to quantify the average number of distinct clones in each sample and observed a significantly increased number of clones in AML compared to MPN or CH (Figure 1E; *FDR p* ≤ 1.37×10^−4^ - 0.026 for all indicated comparisons). The highest number of clones was observed in AML samples with *FLT3* mutations (*FDR p* ≤ 1.37×10^−4^).

**Figure 1.**
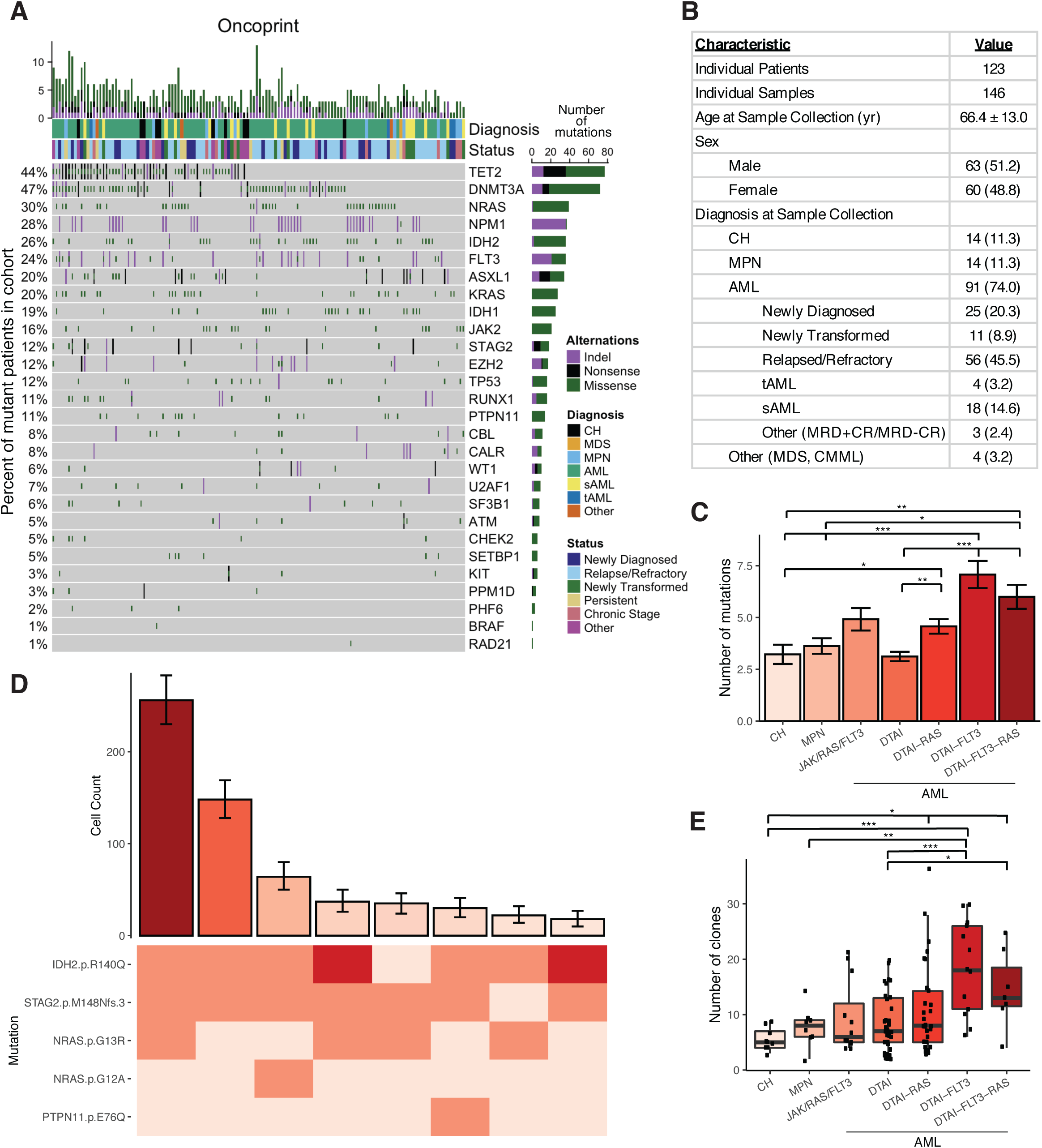
Single cell DNA sequencing of patients with myeloid malignancies. **A)** Oncoprint of patient samples analyzed by single cell DNA sequencing. Each patient is represented with combination of genetic mutations identified by sequencing shown in purple (indel), black (nonsense mutation), and/or green (missense mutation). Percent of samples harboring a specific mutant gene is shown on left of graph with the total number of mutations observed in the cohort for each gene shown on the right of the graph. Diagnosis and disease status at time of sample collection is provided. **B)** Table describing patient cohort characteristics. Standard deviation calculated for mean age of patients at sample collection date. Absolute number of samples denoted with percent of total samples in parentheses. **C)** Bar plot of the number of identified mutations in each sample with samples categorized by cohort. Mean value for each cohort shown by height of bar with standard error measurement (SEM) depicted with error bars. A t-test with false discovery rate (FDR) correction was used to determine statistical significance pairwise between all groups. For clarity, only significant p-values referenced in text are shown. * P < 0.1; ** P < 0.01; ***P < 0.001. **D)** Representative clonal architecture analysis using single cell sequencing. Bar plot depicts the number of cells identified with a given genotype and ranked by decreasing frequency (top panel). Cell counts for each clone is depicted with error bars derived from random resampling analysis. Heatmap shows the genotype consequence of each identified protein coding mutation in the given clone with zygosity (wildtype = light pink, heterozygous = orange, homozygous = red). **E)** Boxplot depicting by cohort the number of unique clones per sample. Each individual sample represented by black circle. A t-test with false discovery rate (FDR) correction was used to determine statistical significance pairwise between all groups. For clarity, only significant p-values referenced in text are shown. FDR p values: * P < 0.1; ** P < 0.01; ***P < 0.001.

We next assessed the diversity of clone size on a per sample basis, calculated as a Shannon diversity index. Clonal diversity significantly increased from CH or MPN to AML (*FDR p* ≤ 0.008; Figure 2A). We observed significantly higher clonal diversity in *RAS* and *FLT3* mutant samples compared to CH/MPN samples and compared to *RAS*/*FLT3*-wildtype AML samples (*FDR p* ≤ 0.039). Despite significant clonal complexity, the majority of AML patients (81.1%; 74/86) had one (75.6%; 65/86) or two (10.5%; 9/86) large clones that accounted for greater than 30% of the cells in the samples. However, the relative size of the largest/dominant clones in different AML subtypes differed significantly. As the clonal diversity increased, the size of the largest clone decreased (*FDR p* ≤ 0.0102 - 0.074 for all indicated comparisons) consistent with the presence of multiple clones with increased fitness (Figure 2B). Despite increased clonal diversity in AML, especially in AML patients with signaling effector mutations, the number of mutations within the largest clone did not significantly differ amongst MPN and AML patients (Extended Figure 3A-B). These data suggest that increased mutational acquisition within the same clone is not the primary driver of clonal dominance. We therefore next investigated whether there were specific genes which when mutated were more likely to be in the dominant clone (Figure 2C, Extended Figure 3C). This analysis revealed gene-specific contributions to clonal expansion, where mutations in *IDH2*, *NPM1* and *JAK2* were nearly always in the dominant clone, while mutations in *FLT3* or *RAS* were more varied, being found only in minor subclones in some patients, and dominant clones in others. The presence of a given mutation in the dominant clone could be inferred from VAF for a subset of frequently mutated genes (Extended Figure 3D), especially *JAK2* which has a known relationship between VAF and clonal dominance in MPN^9,10^.

**Figure 2.**
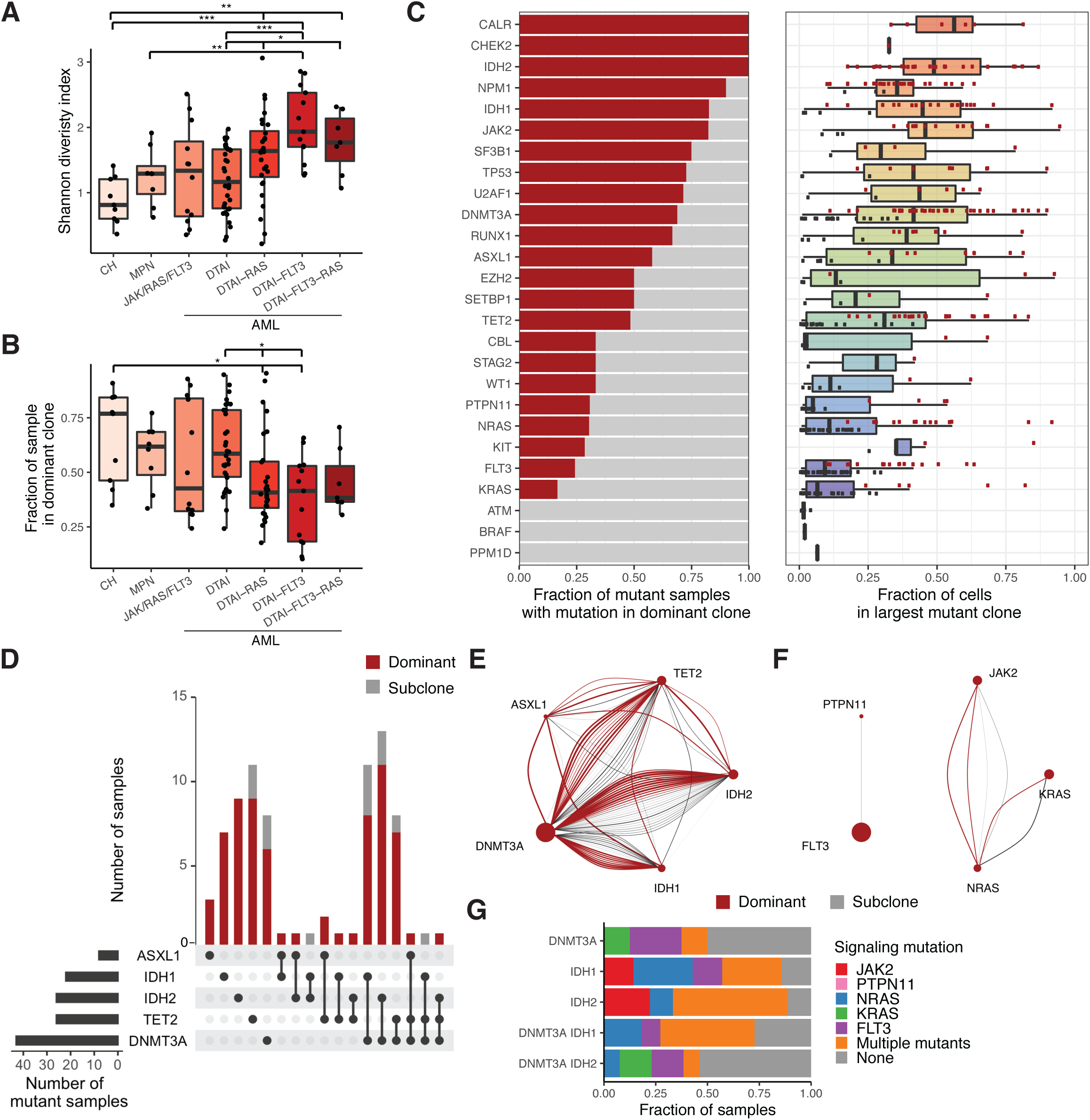
Elucidation of clonal dominance and co-mutation by single cell DNA sequencing. **A)** Box plot depicting by cohort the clonal diversity of each sample calculated by the Shannon diversity index **B)** Box plot depicting by cohort the fraction of total cells in the most dominant clone for each sample. Each individual sample in **A-B** represented by black circle. A t-test with false discovery rate (FDR) correction was used to determine statistical significance pairwise between all groups. For clarity, only significant p-values referenced in text are shown. FDR p values: * P < 0.1; ** P < 0.01; ***P < 0.001. **C)** Bar plot describing the proportion of samples mutant for a given gene where that gene is mutant in the most dominant identified clone (red bar; left panel) with corresponding box plot showing the fraction of cells for a sample in the largest mutant clone (right panel). Each individual sample represented by a circle with samples harboring mutations in the dominant clone (red circle) or a smaller subclone (black). Genes are ranked by decreasing frequency of being identified in the dominant clone within a sample. **D)** Upset plot of co-occurring mutations in samples with more than 1 DTAI variant. Bar graph (top panel) depicts the number of samples with each mutant gene(s) and color of bar annotating whether mutation(s) occur in the dominant clone (red) or subclones (grey). Black circles and connecting line in bottom panel demark the combination of mutations in each corresponding bar plot. **E)** Co-occurrence spectrum of DTAI mutations. Size of circle at each vertex is representative of the number of samples with variants for given gene. Width of connecting arc demarks the relative clone size with color of arc annotating whether co-occurrence is found within dominant clone (red) or subclone (grey). **F)** Network graph of co-occurring signaling mutations. Size of circle at each vertex based on the number of samples with variants for given gene. Width of connecting arc demarks the relative clone size with color of arc annotating whether co-occurrence is found within dominant clone (red) or subclone (grey). Absence of connecting arc denotes no samples with co-occurring mutations found in clinical cohort. **G)** Prevalence of co-occurring signaling mutations in *DNMT3A, IDH1*, and *IDH2* mutant samples. Samples are grouped by presence of *DNMT3A*, *IDH1/2* mutations and fraction of samples mutant for a given signaling mutation is colored accordingly. The absence of a color in the bar plot denotes the absence of a signaling mutation in that sample group.

We next determined mutual inclusivity and exclusivity patterns on a per sample basis and found previously reported mutual exclusivity between *JAK2* and *FLT3* and other reported mutational inverse correlations^3^ (Extended Figure 3E). We then investigated mutational co-occurrence and exclusivity at clonal resolution, focusing on AML patients in our cohort. Amongst the 80 AML samples with DTAI mutations, 52.5% of these samples harbored mutations in more than one epigenetic modifier (Figure 2D). Of note, in nearly all cases these mutations were in the same clone, and in 81% of the cases the co-occurring mutations were within the dominant clone suggesting cooperativity between DTAI mutations in AML (Figure 2E). We did not observe the same co-occurrence patterns of DTAI mutations in CH samples, suggesting that early clonal expansion in the setting of CH is commonly mediated by individual mutations in epigenetic regulators (Extended Figure 3F). By contrast, co-occurring signaling mutations such as *RAS* and *FLT3* very rarely occurred within the same clone, and almost never within the dominant clone (Figure 2F). We further identified distinct mutational cooperativity patterns, including in samples with *DNMT3A* and/or *IDH1/2* mutations (Figure 2G). When *JAK2* was mutated in *IDH1/2* mutant samples, it was the only mutant signaling effector. However, in samples with dual *DNMT3A/IDH* mutant samples, *JAK2* mutations co-occurred with *NRAS* mutations within the same clone (n=3). By contrast, *FLT3* mutations were only observed in *IDH2* mutant samples which also harbored a *DNMT3A* mutation, whereas mutant *FLT3* was present in both single *IDH1* mutant and dual *DNMT3A/IDH1* mutant samples. We identified 6 patients that harbored both *DNMT3A* and *IDH1/2* mutations as well as multiple mutations in signaling effectors (Extended Figure 3G). In these cases, a high fraction of cells across the samples had concurrent *DNMT3A* and *IDH* mutations, whereas few clones were identified with co-mutated signaling genes (*p* ≤ 0.00326), with the exception of a sample with co-mutated *NRAS G13R* and *KRAS G60V* within the same clone. While *NRAS G13R* has been previously identified as an oncogenic mutation, mutations in *KRAS G60V* have not been characterized as being truly oncogenic and may represent a hypomorphic or non-functional allele which does not contribute to transformation and clonal dominance^11^.

Single cell mutation data and consolidated clone architecture provided us an opportunity to delineate the sequence of somatic genetic events during myeloid transformation, and to map these events on clonal expansion. While there are multiple algorithms to model genetic trajectory within a given sample, many do not take into account mutation zygosity (Extended Figure 4A), and therefore necessitated the implementation of an approach that integrated zygosity. We adapted a Markov decision process with reinforcement learning to generate genetic distance networks and trajectories based on the mutations observed in each sample. Using these decision processes, we identified possible trajectories that began with non-mutant wildtype (WT) cells and progressed through both observed and unobserved states in order to maximize the fraction of cells represented in each trajectory. Through this approach, we could determine the optimal initiating mutation and subsequent genetic trajectory based on the path with the highest fraction of observed cells (Figure 3A-B). CH samples displayed oligoclonality and clonal outgrowth of distinct clones with 1-2 mutations (Figure 3A). In AML we observed complex evolutionary trajectories, with clear evidence of clonal dominance and subsequent subclonal propagation (Figure 3B).

**Figure 3.**
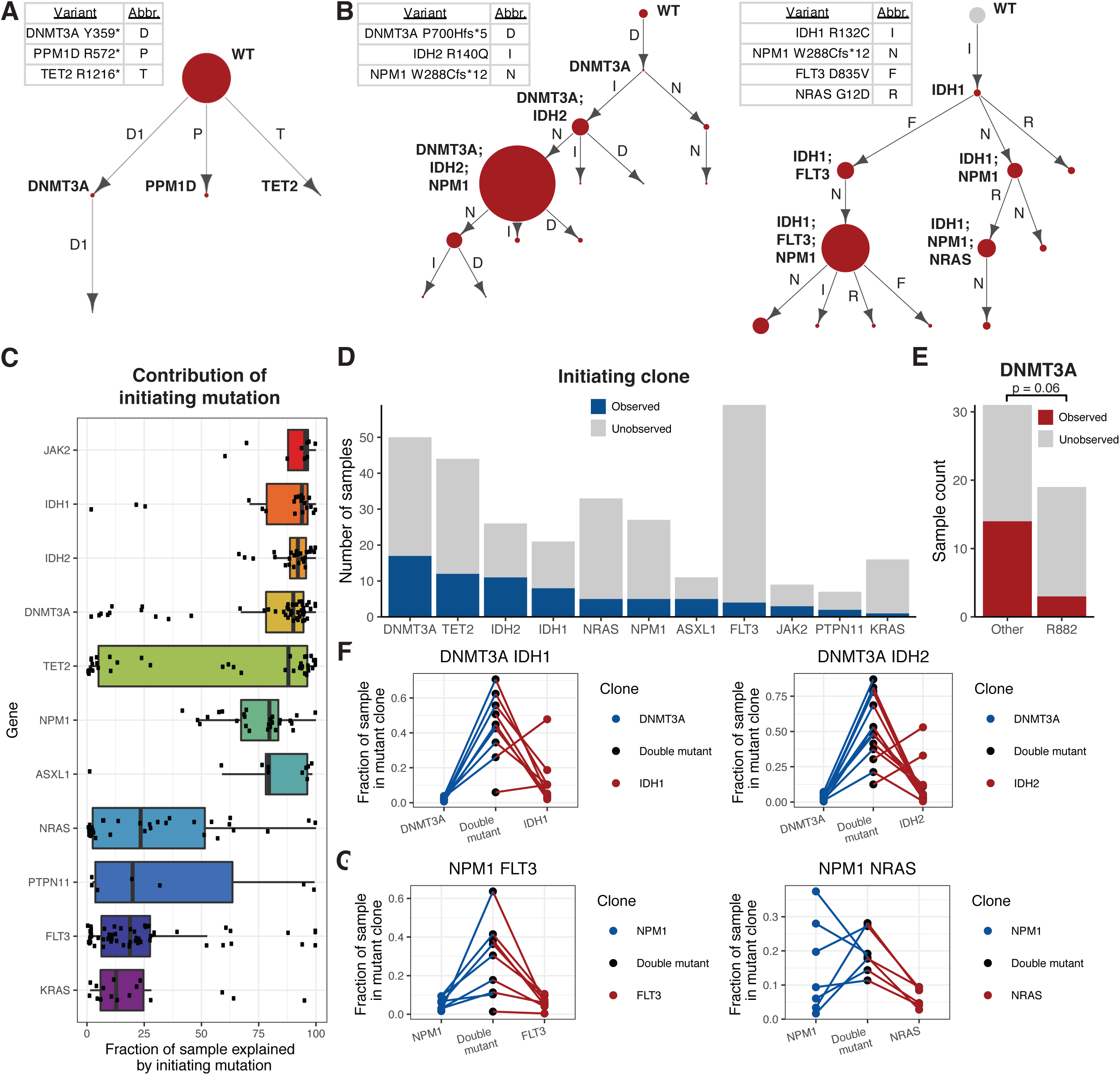
Identification of initiating mutations and clonal expansion through assessing optimal genetic trajectories. **A-B)** Representative optimal genetic trajectories from samples in the CH (**A**) and DTAI cohorts (**B**). Mutant variants for each sample are labeled with abbreviations denoted in corresponding table. Edges are labeled with the mutation acquired. Size of circle denotes relative clone size in the sample with observed clones in red and unobserved clones in grey (with fixed size). Relatively dominant clones are annotated with corresponding acquired mutations. Sample MSK7557 is representative of a CH sample with oligoclonality (**A**). MSK90030 is representative of a sample with more than 1 mutant epigenetic modifier gene (**B**; left panel). MSK87970 is representative of a sample with co-occurring *FLT3, NPM1^c^*, and *RAS* mutations (**B**; right panel) **C)** Fraction of each sample that can be recapitulated when the genetic trajectory starts with a given mutation. Select genes shown are ranked based on mean fraction of samples explained by the genetic trajectories. Each black square denotes a mutant sample from the DTAI cohort with standard deviation denoted. **D**) Number of samples where a single-mutant clone for a given gene is present. Dark blue denotes total number of mutant samples where single-mutant clone is present for a given gene and grey represents mutant samples where single-mutant clone is unobserved. Select genes shown and ordered by decreasing number of samples with present mutant clone from left to right. **E)** Number of *DNMT3A* mutant samples where single-mutant clone is present (red) or absent (grey) with samples categorized by R882 hotspot mutations or other *DNMT3A* mutations. Fisher’s exact test was used to determine statistical significance (*p* ≤ 0.06). **F-G)** Mutant clone sizes (fraction of sample) of single and double mutant clones in *DNMT3A/IDH1* (**F; left panel**) and *DNMT3A/IDH2* (**F; right panel**), *FLT3/NPM1^c^* (**G; left panel**) and *RAS/NPM1^c^* (**G; right panel**) mutant samples. Single-mutant clones and double mutant clones from an individual sample are connected by line between circles in each group. Lines and single-mutant circle are colored are depicted in corresponding color (Figure 3F; *DNMT3A*:blue; *IDH1/IDH2*:red and Figure 3G; *NPM1^c^*:blue; *RAS/FLT3*:red). Double mutant clones are depicted in black.

We next queried samples with DTAI mutations in order to delineate which mutation was the disease-initiating event by assessing the fraction of the clonal architecture possibly explained by a potential initiating mutation (Figure 3C). As expected, the majority of potential states could be reconstructed when the initiating mutation occurred within the epigenetic modifier genes *DNMT3A* and *IDH1/2*. Conversely, very little of the clonal trajectory could be formed if the first mutation were to occur in a signaling gene such as *NRAS* or *FLT3*, with the exception of *JAK2* given its presence as a CH gene and as a disease-defining mutation in MPNs^12–15^. This observation was highly correlated to the computed VAF from scDNA sequencing (Spearman’s correlation coefficient = 0.93; *p* ≤ 2.2 × 10^−16^; Extended Figure 4B). The notable exception was *TET2*, such that it could serve as the disease initiating mutation in some cases and as a mutation which is acquired during clonal progression in other cases. These data are consistent with previous studies in MPN and in post-MPN AML suggesting that *TET2* mutations can be acquired at different times during myeloid transformation and argue for a context-specific effect of *TET2* loss-of-function during myeloid transformation and clonal evolution^16,17^. We next investigated whether mutations in specific genes were observed as initiating, single-mutant clones in our single cell sequencing data, and found that single-mutant clones with a DTAI mutation were commonly identified, confirming these as likely clone-initiating mutations (Figure 3D). However, we observed significant differences in specific alleles within *DNMT3A*, such that *DNMT3A* hotspot R882 mutant-only clones (15.79%; 3/19 mutant samples) were much less frequently detected (*p* < 0.06) than monoallelic clones of all other *DNMT3A* mutants (45.16%; 14/31 mutant samples; Figure 3E). These data suggest that *DNMT3A*-*R882* mutations are either less commonly observed as disease initiating mutations and/or are more likely to acquire additional mutations and undergo rapid clonal evolution. In addition, these data are consonant with the relative paucity of *DNMT3A*-*R882* mutations in CH relative to overt AML^18,19^.

While our analyses strongly suggest the initiating mutations frequently occur in DTAI genes, these analyses do not discern the order of mutations during clonal evolution and relative contribution of these mutations to clonal expansion and dominance. We first focused on samples with co-occurring *DNMT3A*/*IDH1* and *DNMT3A*/*IDH2* mutations to assess mutation order and contribution to clonal dominance in AML patients. For the majority of samples (n=19/23), we observed a significant increase in relative clone size upon the acquisition of the second mutation regardless of whether the clone began with an initiating *DNMT3A* mutation (*IDH1 p* ≤ 0.00023; *IDH2 p* ≤ 2.16 × 10^−6^; Figure 3F), an *IDH1* mutation (*p* ≤ 0.0016), or an *IDH2* mutation (*p* ≤ 1.37 × 10^−5^). We repeated this analysis on *NPM1^c^* mutant samples with co-occurring *FLT3* and *RAS* mutations and determined the fraction of each sample that consisted of single vs. double mutant clones (Figure 3G). Regardless of the mutational order or complexity, the clone size of *FLT3*/*NPM1^c^* double-mutant clones was significantly higher than *FLT3* (*p* ≤ 0.0097) or *NPM1^c^* (*p* ≤ 0.0089) single-mutant clones. By contrast, we observed less evidence of cooperative clonal dominance for *RAS*/*NPM1^c^* co-mutant clones, such that we observed significant variability in clonal size samples when comparing *NPM1^c^* (*p* ≤ 0.462) or *RAS* (*p* ≤ 0.0009) single-mutant clones to the double mutant state. This finding suggests a higher level of mutational synergy between *NPM1^c^* and *FLT3* mutations compared to *NPM1^c^* and *RAS* mutations, and suggest that there is differential contribution of different mutational co-occurrences to clonal dominance in AML, even for commonly observed co-mutant genotypes seen in bulk sequencing.

Recent single cell RNA-sequencing work has demonstrated that leukemia can exist along a continuum of differentiation states and as such, we sought to determine if the genetic clonal architecture we identified above was associated with particular differentiation states^20,21^. We performed simultaneous single cell DNA molecular profiling and cell surface protein expression analysis on AML samples (n = 17) using a DNA + protein single cell sequencing adaptation of the Tapestri platform. We clustered cells from each sample based on immunophenotype and projected DNA genotyping data to identify any association between genotype and immunophenotype (Extended Figure 5A). As expected, we observed significant differences in cell surface protein expression between WT and mutant cells, with WT cells expressing high levels of CD3 (*p* ≤ 2.2 × 10^−16^) and low levels of CD34 (*p* ≤ 2.2 × 10^−16^) compared to mutant cells (Extended Figure 5B). We then compared cell surface protein expression levels in all cells mutant for a given gene across samples (Figure 4A). We found that *TET2* (p ≤ 1.38 × 10^−8^)*, RUNX1* (*p* ≤ 9.48 × 10 ^−13^), *IDH1* (*p* ≤ 2.2 × 10^−16^), and *JAK2* (*p* ≤ 2.2 × 10^−16^) mutant cells were enriched for high CD34 surface expression while cells with mutations in the MAPK/ERK signaling pathway (*NRAS p* ≤ 0.04; *KRAS p* ≤ 2.2 × 10^−16^ and *PTPN11*; *p* ≤ 2.2 × 10^−16^) had higher expression of CD11b compared to other mutant genes. Moreover, we observed that *NPM1^c^* mutant cells harbored lower expression of CD34 compared to all other mutant cells (*p* ≤ 2.2 × 10^−16^), consistent with previous flow cytometric data^22^. Given the combinatorial mutations we identified in our clonal architecture analysis, we next sought to determine how immunophenotype differed across cells harboring multiple mutations. We compared pairwise combinations of *DNMT3A*, *IDH1*, *IDH2*, *FLT3*, *NRAS* and *KRAS*, for their enrichment of different immunophenotypes across all 17 patients (Figure 4B). We observed that the high CD34 expression seen in *IDH1* mutant cells decreased upon the acquisition of signaling effector mutations (*p* ≤ 2.2 × 10^−16^) and concurrently CD11b expression specifically increased in *IDH1/RAS* co-mutant subclones (*p* ≤ 2.2 × 10^−16^).

**Figure 4.**
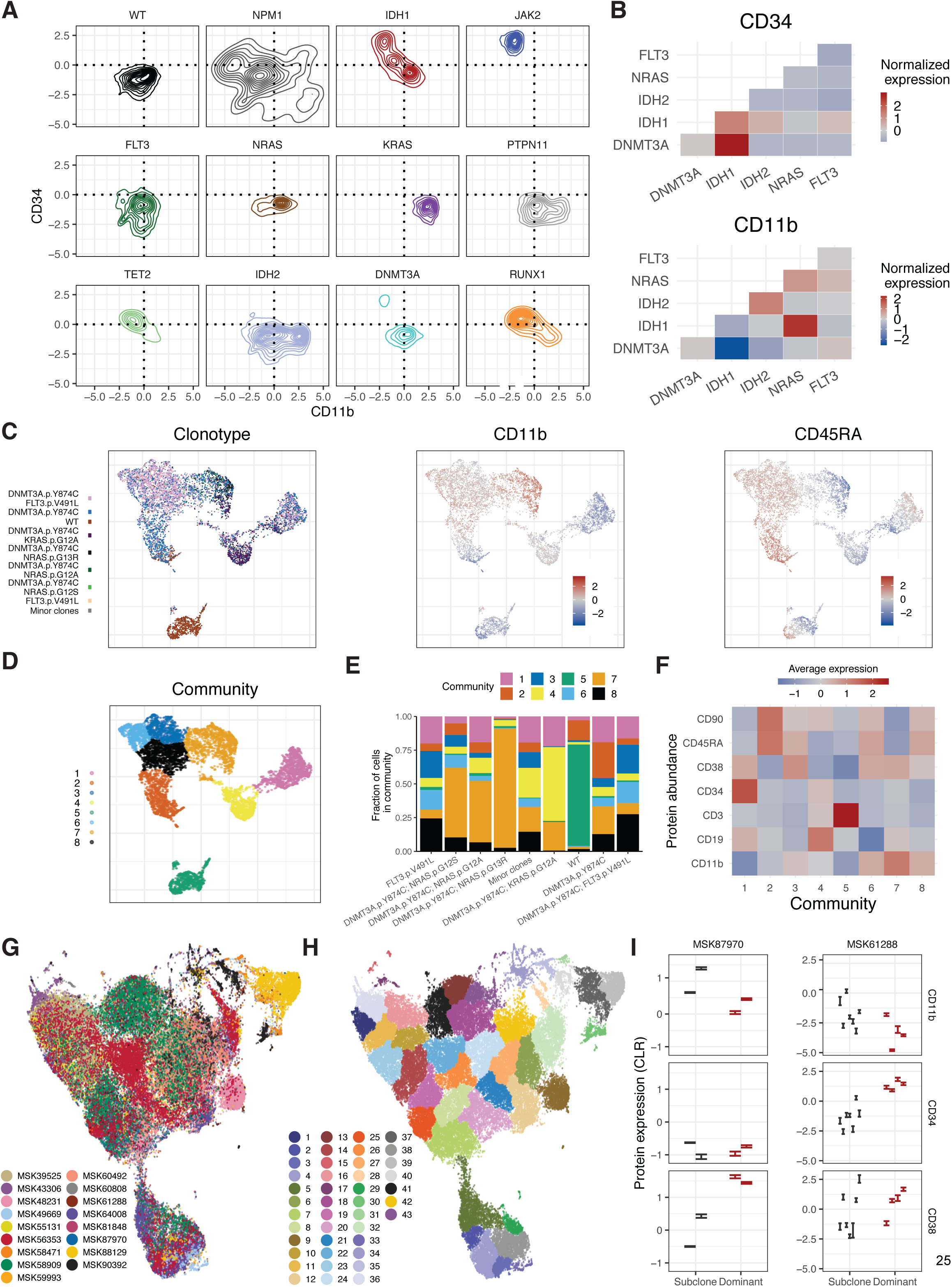
Simultaneous single cell DNA and cell surface protein expression sequencing. **A)** Differential expression of CD34 and CD11b by cells mutant for select genes. Protein expression was normalized by centered log ratio (CLR) transformation. Density plots denotes the frequency of cells with similar protein expression patterns. Origin of x,y-axes denoted with black dashed line. **B)** Immunophenotype changes based on co-occurring mutations in clones. Heatmap of normalized protein expression of CD34 (top panel) and CD11b (bottom panel) in *DNMT3A* and *IDH1/2* single-mutant clones vs. *DNMT3A* and *IDH1/2* mutant clones with co-occurring *NRAS* or *FLT3* mutations. High protein expression depicted in red and low protein expression depicted in blue. **C)** Cell surface expression changes based on genotype of clones. UMAP plot for representative sample MSK56353 clustered by immunophenotype with corresponding clone identity from Extended Figure 5C depicted (top panel) in overlaid colors in left panel. Heatmaps of relative cell surface protein expression overlaid onto UMAP plot for CD11b (middle panel) and CD45RA (right panel) with high relative expression in red and low relative expression in blue. **D)** UMAP for MSK56353 from Figure 4C depicts communities of cells identified by phenoGraph in overlaid colors. **E)** Enrichment of communities based on genotype of clones. Fraction of cells in a given clone clustered in the 8 communities present in MSK56353 is depicted by color of corresponding community. **F)** Immunophenotype differences of communities present in MSK56353. Heatmap depicts the CLR of the 7 cell surface proteins (CD3, CD11b, CD19, CD34, CD38, CD45RA, CD90) with high expression denoted in red and low expression shown by blue. **G**) UMAP plot of samples (n=17) analyzed by DNA+Protein single cell sequencing with cells clustered by cell surface protein expression of 6 antibody targets (CD3, CD11b, CD34, CD38, CD45RA, CD90). Cells from the same sample are denoted with same color. **H)** Neighborhood analysis of all samples. UMAP from Figure 4G depicts communities of cells identified by neighborhood analysis in overlaid colors. **I)** Cell surface protein expression of CD11b, CD34, and CD38 between dominant clone (red) and subclones (black) in a *FLT3-ITD* mutant sample (MSK61288; left panel) and *JAK2* mutant sample (MSK87970; right panel). Each error bar represents a distinct community that is significantly expanded or contracted. A Student’s t-test was used to determine statistical significance. * P < 0.1; ** P < 0.01; ***P < 0.001.

We next characterized how immunophenotype differed across distinct genetic clone populations. We observed expression changes based on the co-occurring mutations contained in specific clones, including increased CD11b expression in the *RAS* mutant subclones compared to *RAS* wildtype subclones (Figure 4C) and reduced CD34 expression of the co-mutant *DNMT3A*/*FLT3* co-mutant clones compared to the *DNMT3A* single-mutant clones (Extended Figure 5C). To summarize combinatorial differences in immunophenotype, we clustered cells into communities and queried changes in immunophenotype on a clonal level^23^ (Figure 4D-F). In one AML patient with an initiating *DNMT3A* mutation and *NRAS, KRAS,* and *FLT3*-mutant subclones, we saw significant differences in community representation in the different genotypes (Figure 4D-E). Specifically, there was an expansion in Community 8 and reduced cells in Community 2 with the acquisition of the *FLT3* mutation in the *DNMT3A* single-mutant clone, concomitant with an increase in CD11b expression and a relative decrease in CD45RA and CD90 surface expression in *FLT3* mutant cells (Figure 4E-F; Extended Figure 5D). We also saw different community enrichment patterns in RAS-mutant clones, however this differed for *NRAS* vs. *KRAS* mutant cells. Cells that acquired mutant *NRAS* showed a marked increase of cells in Community 7 at the expense of all other Communities, which was marked by the highest level of CD11b expression, whereas the acquisition of *KRAS* increased representation in Community 4, associated with high CD19 expression (Extended Figure 5D).

To determine if patterns of immunophenotype changes existed across multiple samples, we merged all samples, clustered cells based on cell surface protein expression, then identified communities of cells using phenoGraph^23^ (Figure 4G-H). Our findings suggest that multiple overlapping immunophenotypic states occur across samples even with divergent genotypes; no community was exclusive to an individual sample and 6 communities were observed in every sample (Community 7, 8, 9, 18, 32, and 42) which were intercorrelated with high expression of either CD90 or CD38 (Extended Figure 6A). We next sought to determine if samples within our cohort showed recurrent community representation alterations between domain and subclones (Extended Figure 6B). We observed significant shifts in communities between the dominant clone and subclones in 8/14 samples, with the remaining 3 samples possessing a single variant and excluded from clonality analysis. In contrast to our findings correlating the increase of CD11b expression to the acquisition of *NRAS* mutations, we also observed an expansion of a community high in CD34 expression (*p* ≤ 2.2 × 10^−16^) and low in CD11b expression (*p* ≤ 2.2 × 10^−16^) in a sample (MSK87970) upon the acquisition of a *FLT3-ITD* mutation (Figure 4I). Furthermore, a *JAK2* mutant sample (MSK61288) showed an expansion of communities with high CD38 and with low CD11b expression in the dominant clone compared to subclones (CD38; *p* ≤ 2.2 × 10^−16^; CD11b; *p* ≤ 2.2 × 10^−16^). These findings suggest divergent clone-specific changes in cell surface expression upon the acquisition of mutant signaling effectors.

## Discussion

The identification of highly frequent, recurrent mutations in epigenetic regulators in CH, and the relative paucity of cases of overt MPN, MDS and AML compared to CH suggests that the rate-limiting step in myeloid transformation is clonal evolution from disease-initiating clones to leukemic clones through stepwise mutational acquisition. Previous studies have used bulk sequencing analyses to predict important features of clonal evolution^3,24–26^; however, the molecular sequence of events which drive myeloid transformation have not been dissected at a single-cell, clonal level. Here we use single cell molecular profiling to precisely map clonal evolution in myeloid malignancies, and to make several important insights into the pathogenesis of myeloid transformation. First, we found that the number of mutations and clonal complexity increases from CH or MPN to AML and continues to evolve as AML clones acquire mutations in signaling effectors such as *RAS* or *FLT3*. Second, we found that specific genes and gene combinations were capable of driving clonal expansion and dominance, such that they were almost always found in the dominant clone. This included co-occurring mutations in epigenetic regulators as well as disease-defining mutations such as *JAK2* and *NPM1^c^*. By contrast, signaling effector mutations were often subclonal, and very rarely co-occurring in the same clone. In addition, when looking at how cooperating alleles influenced the genetic trajectory and clonal trajectory in AML, we observed significant differences in how mutational combinations contributed to clonal dominance such that specific co-occurring disease alleles (e.g. *NPM1^c^* + *FLT3-ITD* or *DNMT3A* + *IDH2*) were associated with clonal dominance and other mutational combinations (*NPM1^c^* + *RAS*) did not promote clonal expansion. Lastly, we identified significant changes in cell surface protein expression driven by genotypes using simultaneous single cell mutational profiling and immunophenotyping. Signaling effector mutations were particularly notable for altering cell surface protein expression, with acquisition of MAPK/ERK pathway mutations leading to increased CD11b expression.

While the ability to profile mutational spectra and immunophenotype at single cell resolution is powerful, our analyses have limitations which will be attenuated as single cell DNA sequencing increases in throughput (to 100,000-1,000,000 cells/sample). Technical advances with unique molecular identifiers will help resolve further analysis into mutation order and architecture through accurate assessment of very small clones. We analyzed each patient at a single timepoint, and analysis of serial samples will delineate how clonal evolution changes during disease progression and/or in response to anti-leukemic therapies. Moreover, our analyses of clones and trajectories assume that the clones expand and contract without inter-clonal interactions. There is growing evidence that mutant clones can interact with other existing clones and could potentially affect the growth and fitness of each other. The observation that the majority of AML samples are comprised of 1-2 dominant clones adds an important layer to this possibility, such that dominant clone(s) may outcompete minor clones through increased proliferation/self-renewal or through active, cell non-autonomous suppression of less fit minor clones as has been shown in murine models of CML where mutant cells can suppress the fitness of wild-type cells^27^. Taken together, these data suggest that patients with myeloid malignancies manifest as a complex ecosystem of clones which evolves over time, and that single cell sequencing gives a glimpse into this milieu not seen with conventional bulk sequencing. In summary, our studies of clonal architecture at a single cell level provide us novel insights into the pathogenesis of myeloid transformation and give us new insights into how clonal complexity contributes to disease progression. Similar studies across different pre-malignant and malignant contexts will give new insights into how malignancy initiate and progresses and will lead to new therapeutic insights aimed at intercepting clonal evolution and/or targeting cancer as a multi-clonal disease.

## Online Methods

### Reagents

All antibodies were purchased from Biolegend. These studies used the following antibodies: FITC-CD3 (clone UCHT1) FITC-CD19 (clone HIB19) and FITC-CD56 (clone HCD56). Human TruStain FcX was also purchased from Biolegend. The DNA+Protein oligo-conjugated antibodies were produced and purchased from Biolegend. The antibodies in the conjugate pool were the following: CD3 (clone SK7), CD11b (clone RCRF44), CD19 (clone HIB19), CD34 (clone 581), CD38 (clone HIT2), CD45RA (clone HI100), and CD90 (clone 5E10). Antibody conjugates were pooled in equimolar ratios. All Tapestri related reagents were included as part of a Custom Single Cell DNA sequencing kit purchased from Mission Bio, Inc. The Custom amplicon panel used in these studies covers 109 amplicons over 31 genes previously found to be frequently mutated in human myelodysplastic syndromes (MDS), myeloproliferative neoplasms (MPN), and acute myeloid leukemia (AML) (Extended Table 1)^1–3^.

### Patient Samples

Patients with myeloid neoplasms or acute myeloid leukemia between 2014 and 2019 were studied. Informed consent was obtained from patients according to protocols approved by the institutional IRBs and in accordance with the Declaration of Helsinki. This study was approved by MSKCC Institutional Review Board (protocol #15-017) and Thomas Jefferson University (TJU) Institutional Review Board (protocol# 17D.083). Diagnosis and disease status was confirmed and assigned according to World Health Organization (WHO) classification criteria^4^. Patient characteristics are summarized in Extended Table 2 and Figure 1B. Bone marrow from healthy individuals was obtained with informed consent according to procedures approved by the institutional review boards Memorial Sloan Kettering Cancer Center and Hospital for Special Surgery. Patient samples were collected and processed by the MSKCC Human Oncology Tissue Bank (HOTB) or TJU Heme Malignancy Repository. Mononuclear cells were obtained by centrifugation on Ficoll from peripheral blood or bone marrow and viably frozen. Patient samples from MSKCC underwent high-throughput genetic sequencing with a targeted deep sequencing assay of 685 genes (HemePACT) or by an NGS platform panel composed of 49 genes recurrently mutated in myeloid disorders (RainDance Technologies ThunderBolts Myeloid Panel). Single point variants were called using Mutect and short insertions and deletions using Pindel as described previously, comparing samples to a sample representing a pool of normal samples^5^. Mutations were excluded if found to be present in at least one database of known non-somatic variants (dbSNP and 1000 genomes) and absent from COSMIC. Samples with non-excluded mutations with variant allele frequency >2% were classified as clonal hematopoiesis. Samples were selected based on mutation coverage by the Mission Bio Custom amplicon panel, variant allele frequencies of all covered mutations (>5% VAF for each gene covered on panel), and number of cells collected (>5 × 10^6^ cells) per frozen aliquot. Specifically, samples were prioritized if they harbored 1) more than one mutation in epigenetic modifier genes *DNMT3A*, *TET2*, *ASXL1*, or *IDH1/2*, 2) a *NPM1* mutation, 3) mutations in *NRAS*, *KRAS*, and/or 4) mutations in *FLT3* (either internal tandem duplication (ITD) or tyrosine kinase domain (TKD) mutations).

### Single cell DNA sequencing library preparation and sequencing

Patient samples were thawed and washed with PBS supplemented with 1% BSA (FACS buffer). Cells were incubated with TruStain FcX (Biolegend) for 15 min at 4°C then stained with FITC-conjugated antibodies against human CD3, CD19, and CD56 (NCAM) for 15 min at 4°C. Cells were then washed and resuspended in FACS buffer with DAPI and sorted to isolate viable (DAPI^−^) CD3^−^/CD19^−^/CD56^−^ (FITC^−^) cells using a Sony SH800 Cell Sorter. Cells were resuspended in Tapestri cell buffer and quantified using a Countess cell counter (Invitrogen). Single cells (3-4,000 cells/μL) were encapsulated using a Tapestri microfluidics cartridge, lysed, and barcoded^6^. Barcoded samples were then subjected to targeted PCR amplification of a custom 109 amplicons covering 31 genes known to be involved in hematologic malignancies (AML/MPN/MDS; Extended Table 1). PCR products were removed from individual droplets, purified with Ampure XP beads (Beckman Coulter), and used as a template for PCR to incorporate Illumina i5/i7 indices. PCR products were purified a second time, quantified via an Agilent Bioanalyzer and pooled to be sequenced. Library pools were sequenced on an Illumina NovaSeq by the MSKCC Integrated Genomics Core.

### Single cell DNA & Protein sequencing library preparation and sequencing

Patient samples were thawed, washed with FACS buffer, and quantified using a Countess cell counter. Cells (1.0-4.0 × 10^6^ viable cells) were then resuspended in DPBS (Gibco) and incubated with TruStain FcX, Dextran Sulfate (100 μg/mL; Research Products International), and 1X Tapestri staining buffer for 3 minutes at room temperature. The pool of 7 oligo-conjugated antibodies (CD3, CD11b, CD19, CD34, CD38, CD45RA, CD90) was then added and incubated for 30 minutes at room temperature. Cells were then washed multiple times with DPBS supplemented with 5% fetal bovine serum (FBS; Gibco) followed by resuspension of the cells in Tapestri cell buffer, requantification, and loading of the cells into a Tapestri microfluidics cartridge. Single cells were encapsulated, lysed, barcoded as above with the exception of adding an additional forward primer mix (3 μM each) for the antibody tags prior to barcoding. DNA PCR products were then isolated from individual droplets and purified with Ampure XP beads. The DNA PCR products were then used as a PCR template for library generation as above and repurified using Ampure XP beads. Protein PCR products (supernatant from Ampure XP bead incubation) were incubated with Tapestri pullout oligo (5 μM) at 96°C for 5 minutes followed by incubation on ice for 5 minutes. Protein PCR products were then purified using Steptavidin C1 beads (Invitrogen) and beads were used as a PCR template for the incorporation of i5/i7 Illumina indices followed by purification using Ampure XP beads. All libraries, both DNA and Protein, were quantified using an Agilent Bioanalyzer and pooled for sequencing on an Illumina NovaSeq by the MSKCC Integrated Genomics Core.

### Data Analysis

#### Data processing

FASTQ files for single cell DNA libraries were analyzed through the Tapestri Pipeline using Bluebee’s high performance genomics platform. Briefly, this pipeline trims adaptor sequences, aligns reads to the human genome (hg19), assigns sequence reads to cell barcodes, and performs genotype calling with GATK. Data is then consolidated into a multiple sample VCF file and output as a loom file for subsequent processing. Initial steps for filtering low quality genotypes or cells was performed in Tapestri Insights and R, where the minimum variant quality score was set to 30 with a minimum of 10 reads per variant per cell. We further removed variants present in <50% of cells and removed cells in which <50% of potential variants reported informative genotypes. Data was output from Tapestri Insights and subsequent filtering was performed in R. For DNA analysis on the DNA+Protein platform, we used the Tapestri Pipeline on Bluebee as described above. For the protein analysis, custom scripts in R were used by Mission Bio to enumerate the number of reads per antibody per cell. Subsequent normalization was performed using the tapestri package in R. Samples were included if they harbored 1 or more protein encoding, non-synonymous/insertion/deletion variants and more than 100 cells with definitive genotype for all protein coding variants within the sample. We next sought to define genetic clones, which we identified as cells that possessed identical genotype calls for the protein encoding variants of interest. In order to focus our analyses on reproducible clones, we performed a bootstrapping analysis over 10,000 samplings to calculate 95% confidence intervals for the presence of each clone. Clonal analyses in Figure 1C and onward focus on clones where the lower 95% confidence interval > 10 cells. We further excluded rare variants which were only identified in clones that did not pass this threshold. Samples were included in clonal analyses if they encoded >2 protein encoding variants and >2 clones. Flowchart of sample inclusion can be found in Extended Figure 2, and patient characteristics can be found in Extended Table 2. Dominant clones as referred to in the text were defined as the largest mutant clone in the sample, excluding cells which were wild type for all variants of interest.

#### Genetic trajectory analysis

For the genetic trajectory analysis constructed in Figure 3, we implemented a markov decision process with reinforcement learning. Generally, this allowed us to model the optimal track of mutation acquisition if a cell were to acquire one mutation at a time and not revert that mutation to a wild type state. Technically, for a given sample, we first constructed a reward matrix by enumerating all possible clones given the number of mutations present in a sample, and the maximum zygosity for a given mutant (i.e., if we did not observe a homozygous state for a mutant, it was not considered in the reward matrix). After construction of the reward matrix, we set permissible decision processes with a value of 0, and impermissible decision processes with a value of -1 (i.e. decisions where a mutant was reverted to wildtype or required more than one genetic alteration were penalized). Decisions were considered permissible if a clone was separated by a single genetic event, either a variant changing from wildtype to heterozygous or heterozygous to homozygous. For observed clones, the frequency of the clone (ranging from 0-100% of cells) was used as the value in the reward matrix, while unobserved clones retained a value of 0. The matrix was then converted to long form and state transitions between clones were associated with the action/mutation causative to that state change. This was then used as input to the ReinforcmentLearning package in R to generate a Q matrix through the experience replay algorithm^7^. Custom scripts in R were used to navigate this Q matrix to determine optimal trajectory from the wildtype clone.

#### Statistical analysis

Statistical significance was evaluated using a Student’s T-test and Fisher’s exact test where indicated. Multiple test correction was implemented using the Benjamini-Hochberg/FDR approach as indicated. Shannon diversity index was assessed using the diversity function in the vegan package in R^8^. Genetic co-occurrence analysis was performed using the cooccur package in R. UMAP clustering was performed using the R package umap, with default paramaters^9,10^. Subsequent community analysis was performed using phenograph implemented with the Rphenograph package^11,12^. The perplexity factor K was set to 50.

#### Plotting and graphical representations

All barplots, boxplots, heatmaps and scatterplots were produced using the ggplot2 package in R^13^. Error bars depict standard error of measure. Boxplots are depicted in Tukey’s style with boxes representing the median and quartile range, with whiskers representing +/− 1.5x the IQR. The oncoprint presented in Figure 1A was produced using the ComplexHeatmap package in R^14^. Upset plots shown in Figure 2D and Extended Figure 3F were produced using the UpsetR package^15^. Network plots in Figure 2E, F and Figure 3A-C were produced with the igraph package in R^16^. UMAP data was plotted using the ggplot2 package. Other packages used in data processing include tidyr, dplyr, RColorbrewer, pals, and cowplot.

#### Data Availability

All scripts and processed data files are available at https://github.com/bowmanr/scDNA_myeloid. Raw data are available upon request from the authors and are being uploaded to dbGAP prior to final publication.

## Acknowledgements

We acknowledge the use of the MSKCC Integrated Genomics Core for all library sequencing which is funded by the MSKCC Support Grant NIH P30 CA008748. L.A.M is supported by a Career Development Program Fellowship of the Leukemia and Lymphoma Society (5479-19). R.L.B. is supported by the Sohn Foundation Fellowship of the Damon Runyon Cancer Research Foundation (DRG 22-17). A.D.V. is supported by the William Raveis Charitable Fund Fellowship of the Damon Runyon Cancer Research Foundation (DRG 117-15), an EvansMDS Young Investigator grant from the Edward P. Evans Foundation, and a National Cancer Institute career development grant K08 CA215317. This work is supported by grants to S.E.M including National Cancer Institute R37 CA226433, Conquer Cancer Now Award from the Concern Foundation, and Sidney Kimmel Cancer Center (SKCC) Support Grant NIH P30 CA056036. This work was supported by grants to R.L.L. including a Cycle For Survival Innovation Grant, National Cancer Institute R35 CA197594, National Cancer Institute R01 CA173636, and a SCOR grant from the Leukemia and Lymphoma Society.

## Author Contributions

L.A.M., R.L.B., A.D.V., and R.L.L. conceptualized studies. L.A.M., R.L.B, A.O., R.D.D., P.M., C.A., M.M., S.S., M.S.B., and R.L.L. designed and optimized experimental methodologies and bioinformatic workflow. L.A.M., R.L.B., T.R.M., I.S.C., P.M. and A.O. performed experiments. C.F, M.A.P., K.B., A.Z., A.D.G., M.P.C., S.E.M., R.R., A.D.V., and R.L.L. provided patient associated resources and/or patient samples for use in studies. L.A.M., R.L.B., and R.L.L wrote the original manuscript. L.A.M., R.L.B., S.E.M., and R.L.L assisted with review and editing of the manuscript. R.L.L supervised studies and manuscript preparation.

## Extended Data

**Extended Table 1.**
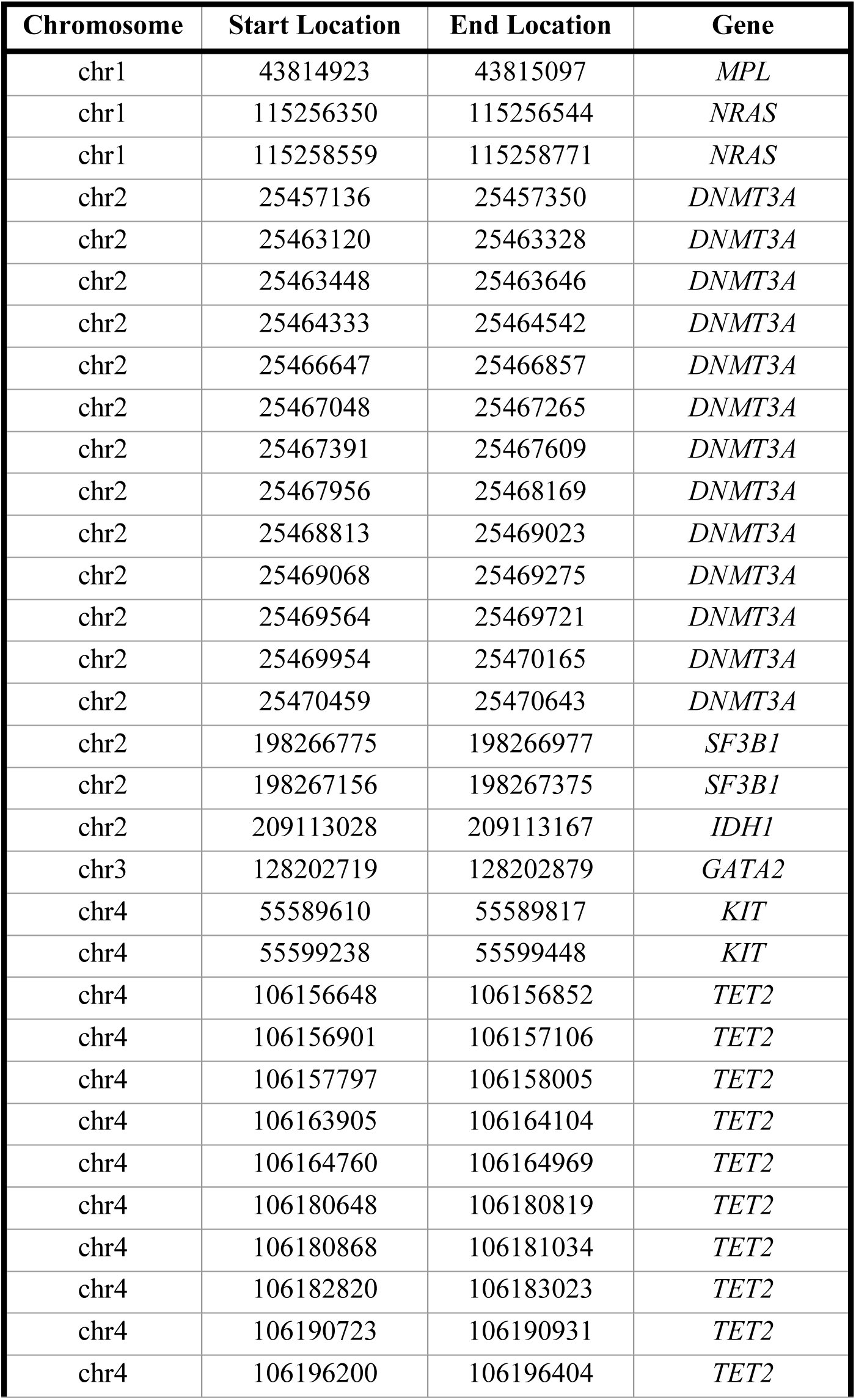
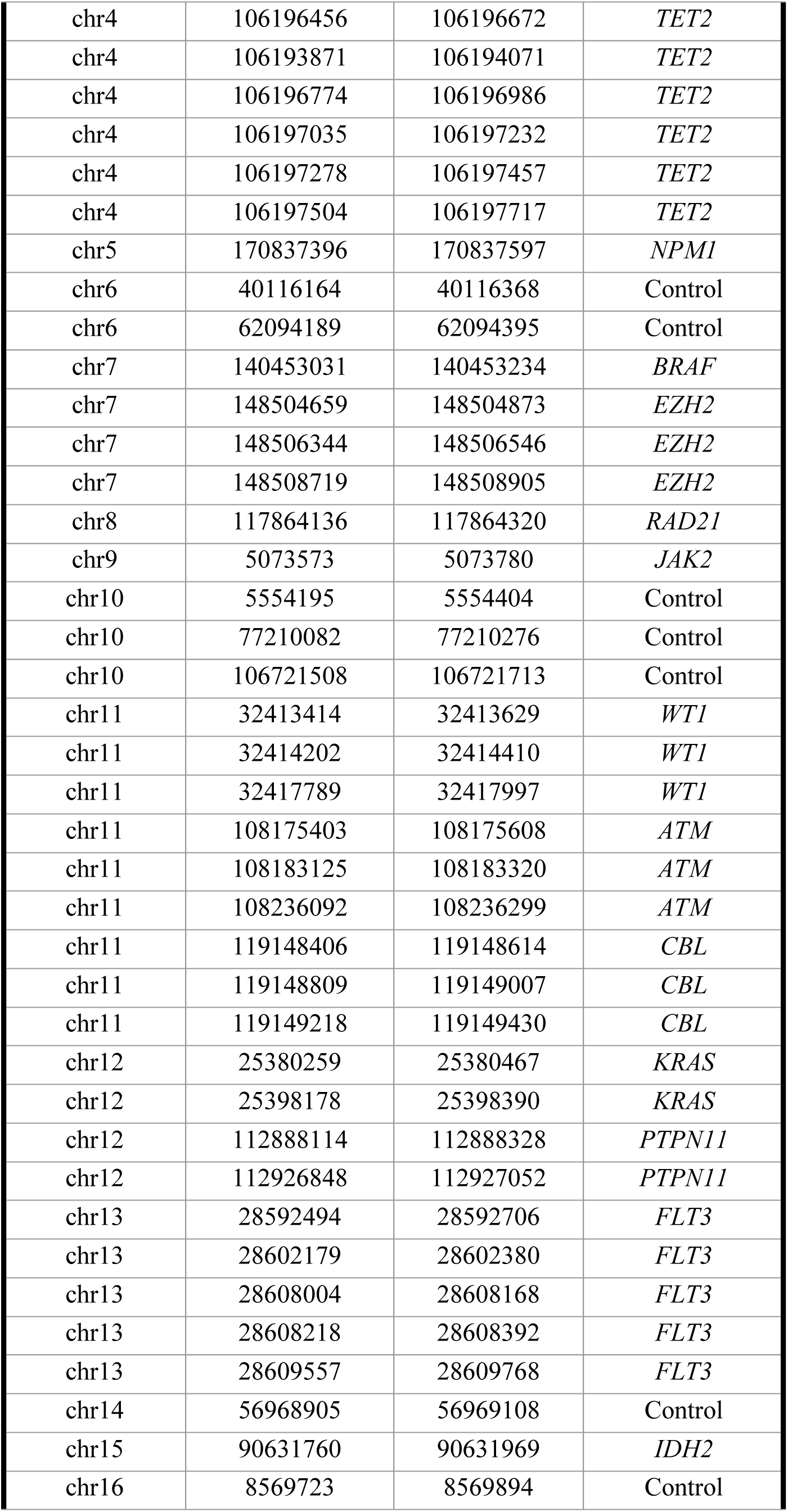
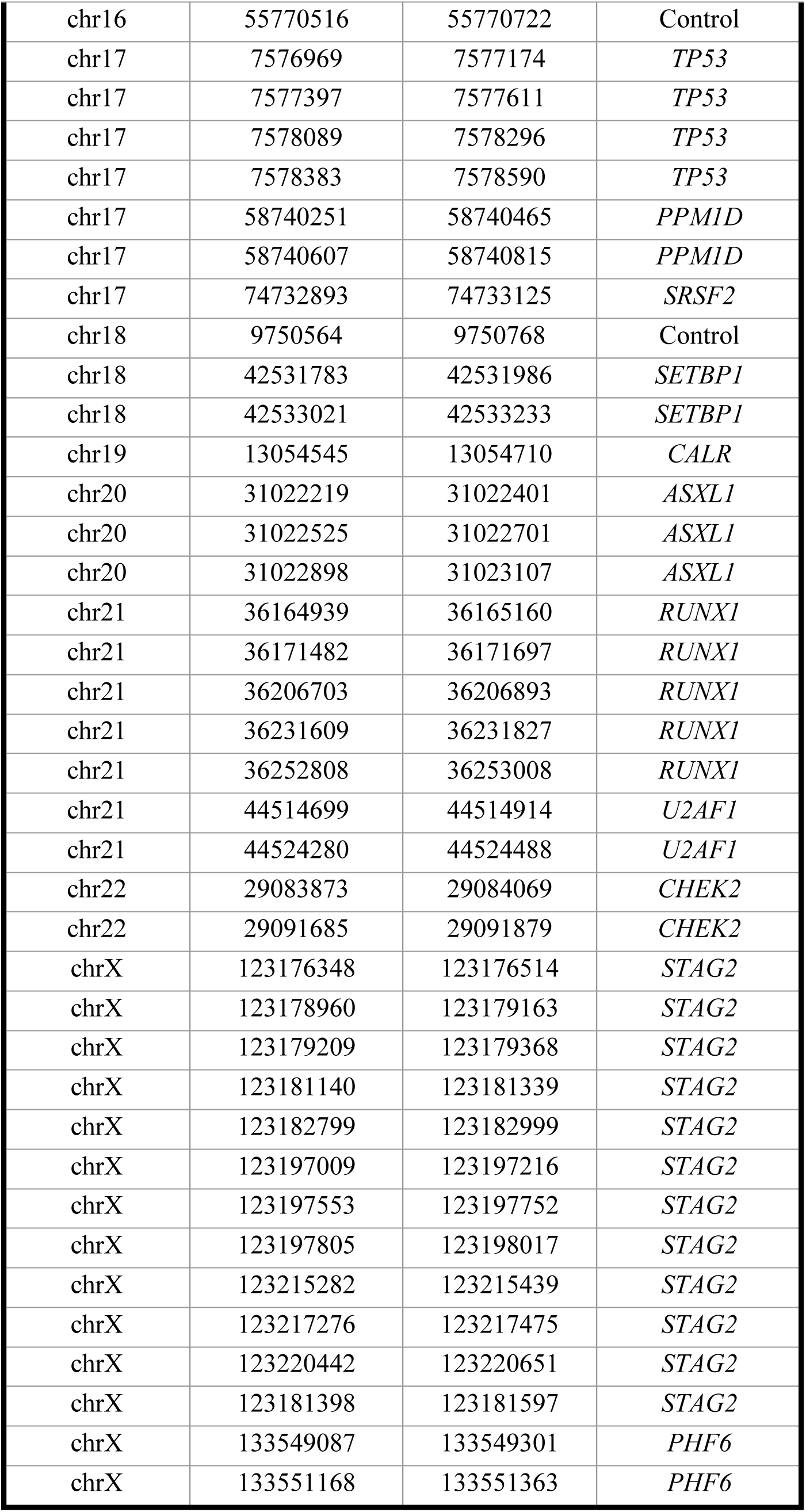
Amplicon Coverage for Custom Mission Bio single cell DNA sequencing panel. Each amplicon is listed with genomic location (hg19) and gene name.

| Chromosome | Start Location | End Location | Gene |
| --- | --- | --- | --- |
| chr1 | 43814923 | 43815097 | *MPL* |
| chr1 | 115256350 | 115256544 | *NRAS* |
| chr1 | 115258559 | 115258771 | *NRAS* |
| chr2 | 25457136 | 25457350 | *DNMT3A* |
| chr2 | 25463120 | 25463328 | *DNMT3A* |
| chr2 | 25463448 | 25463646 | *DNMT3A* |
| chr2 | 25464333 | 25464542 | *DNMT3A* |
| chr2 | 25466647 | 25466857 | *DNMT3A* |
| chr2 | 25467048 | 25467265 | *DNMT3A* |
| chr2 | 25467391 | 25467609 | *DNMT3A* |
| chr2 | 25467956 | 25468169 | *DNMT3A* |
| chr2 | 25468813 | 25469023 | *DNMT3A* |
| chr2 | 25469068 | 25469275 | *DNMT3A* |
| chr2 | 25469564 | 25469721 | *DNMT3A* |
| chr2 | 25469954 | 25470165 | *DNMT3A* |
| chr2 | 25470459 | 25470643 | *DNMT3A* |
| chr2 | 198266775 | 198266977 | *SF3B1* |
| chr2 | 198267156 | 198267375 | *SF3B1* |
| chr2 | 209113028 | 209113167 | *IDH1* |
| chr3 | 128202719 | 128202879 | *GATA2* |
| chr4 | 55589610 | 55589817 | *KIT* |
| chr4 | 55599238 | 55599448 | *KIT* |
| chr4 | 106156648 | 106156852 | *TET2* |
| chr4 | 106156901 | 106157106 | *TET2* |
| chr4 | 106157797 | 106158005 | *TET2* |
| chr4 | 106163905 | 106164104 | *TET2* |
| chr4 | 106164760 | 106164969 | *TET2* |
| chr4 | 106180648 | 106180819 | *TET2* |
| chr4 | 106180868 | 106181034 | *TET2* |
| chr4 | 106182820 | 106183023 | *TET2* |
| chr4 | 106190723 | 106190931 | *TET2* |
| chr4 | 106196200 | 106196404 | *TET2* |
| chr4 | 106196456 | 106196672 | *TET2* |
| chr4 | 106193871 | 106194071 | *TET2* |
| chr4 | 106196774 | 106196986 | *TET2* |
| chr4 | 106197035 | 106197232 | *TET2* |
| chr4 | 106197278 | 106197457 | *TET2* |
| chr4 | 106197504 | 106197717 | *TET2* |
| chr5 | 170837396 | 170837597 | *NPM1* |
| chr6 | 40116164 | 40116368 | Control |
| chr6 | 62094189 | 62094395 | Control |
| chr7 | 140453031 | 140453234 | *BRAF* |
| chr7 | 148504659 | 148504873 | *EZH2* |
| chr7 | 148506344 | 148506546 | *EZH2* |
| chr7 | 148508719 | 148508905 | *EZH2* |
| chr8 | 117864136 | 117864320 | *RAD21* |
| chr9 | 5073573 | 5073780 | *JAK2* |
| chr10 | 5554195 | 5554404 | Control |
| chr10 | 77210082 | 77210276 | Control |
| chr10 | 106721508 | 106721713 | Control |
| chr11 | 32413414 | 32413629 | *WT1* |
| chr11 | 32414202 | 32414410 | *WT1* |
| chr11 | 32417789 | 32417997 | *WT1* |
| chr11 | 108175403 | 108175608 | *ATM* |
| chr11 | 108183125 | 108183320 | *ATM* |
| chr11 | 108236092 | 108236299 | *ATM* |
| chr11 | 119148406 | 119148614 | *CBL* |
| chr11 | 119148809 | 119149007 | *CBL* |
| chr11 | 119149218 | 119149430 | *CBL* |
| chr12 | 25380259 | 25380467 | *KRAS* |
| chr12 | 25398178 | 25398390 | *KRAS* |
| chr12 | 112888114 | 112888328 | *PTPN11* |
| chr12 | 112926848 | 112927052 | *PTPN11* |
| chr13 | 28592494 | 28592706 | *FLT3* |
| chr13 | 28602179 | 28602380 | *FLT3* |
| chr13 | 28608004 | 28608168 | *FLT3* |
| chr13 | 28608218 | 28608392 | *FLT3* |
| chr13 | 28609557 | 28609768 | *FLT3* |
| chr14 | 56968905 | 56969108 | Control |
| chr15 | 90631760 | 90631969 | *IDH2* |
| chr16 | 8569723 | 8569894 | Control |
| chr16 | 55770516 | 55770722 | Control |
| chr17 | 7576969 | 7577174 | *TP53* |
| chr17 | 7577397 | 7577611 | *TP53* |
| chr17 | 7578089 | 7578296 | *TP53* |
| chr17 | 7578383 | 7578590 | *TP53* |
| chr17 | 58740251 | 58740465 | *PPM1D* |
| chr17 | 58740607 | 58740815 | *PPM1D* |
| chr17 | 74732893 | 74733125 | *SRSF2* |
| chr18 | 9750564 | 9750768 | Control |
| chr18 | 42531783 | 42531986 | *SETBP1* |
| chr18 | 42533021 | 42533233 | *SETBP1* |
| chr19 | 13054545 | 13054710 | *CALR* |
| chr20 | 31022219 | 31022401 | *ASXL1* |
| chr20 | 31022525 | 31022701 | *ASXL1* |
| chr20 | 31022898 | 31023107 | *ASXL1* |
| chr21 | 36164939 | 36165160 | *RUNX1* |
| chr21 | 36171482 | 36171697 | *RUNX1* |
| chr21 | 36206703 | 36206893 | *RUNX1* |
| chr21 | 36231609 | 36231827 | *RUNX1* |
| chr21 | 36252808 | 36253008 | *RUNX1* |
| chr21 | 44514699 | 44514914 | *U2AF1* |
| chr21 | 44524280 | 44524488 | *U2AF1* |
| chr22 | 29083873 | 29084069 | *CHEK2* |
| chr22 | 29091685 | 29091879 | *CHEK2* |
| chrX | 123176348 | 123176514 | *STAG2* |
| chrX | 123178960 | 123179163 | *STAG2* |
| chrX | 123179209 | 123179368 | *STAG2* |
| chrX | 123181140 | 123181339 | *STAG2* |
| chrX | 123182799 | 123182999 | *STAG2* |
| chrX | 123197009 | 123197216 | *STAG2* |
| chrX | 123197553 | 123197752 | *STAG2* |
| chrX | 123197805 | 123198017 | *STAG2* |
| chrX | 123215282 | 123215439 | *STAG2* |
| chrX | 123217276 | 123217475 | *STAG2* |
| chrX | 123220442 | 123220651 | *STAG2* |
| chrX | 123181398 | 123181597 | *STAG2* |
| chrX | 133549087 | 133549301 | *PHF6* |
| chrX | 133551168 | 133551363 | *PHF6* |

**Extended Table 2.**
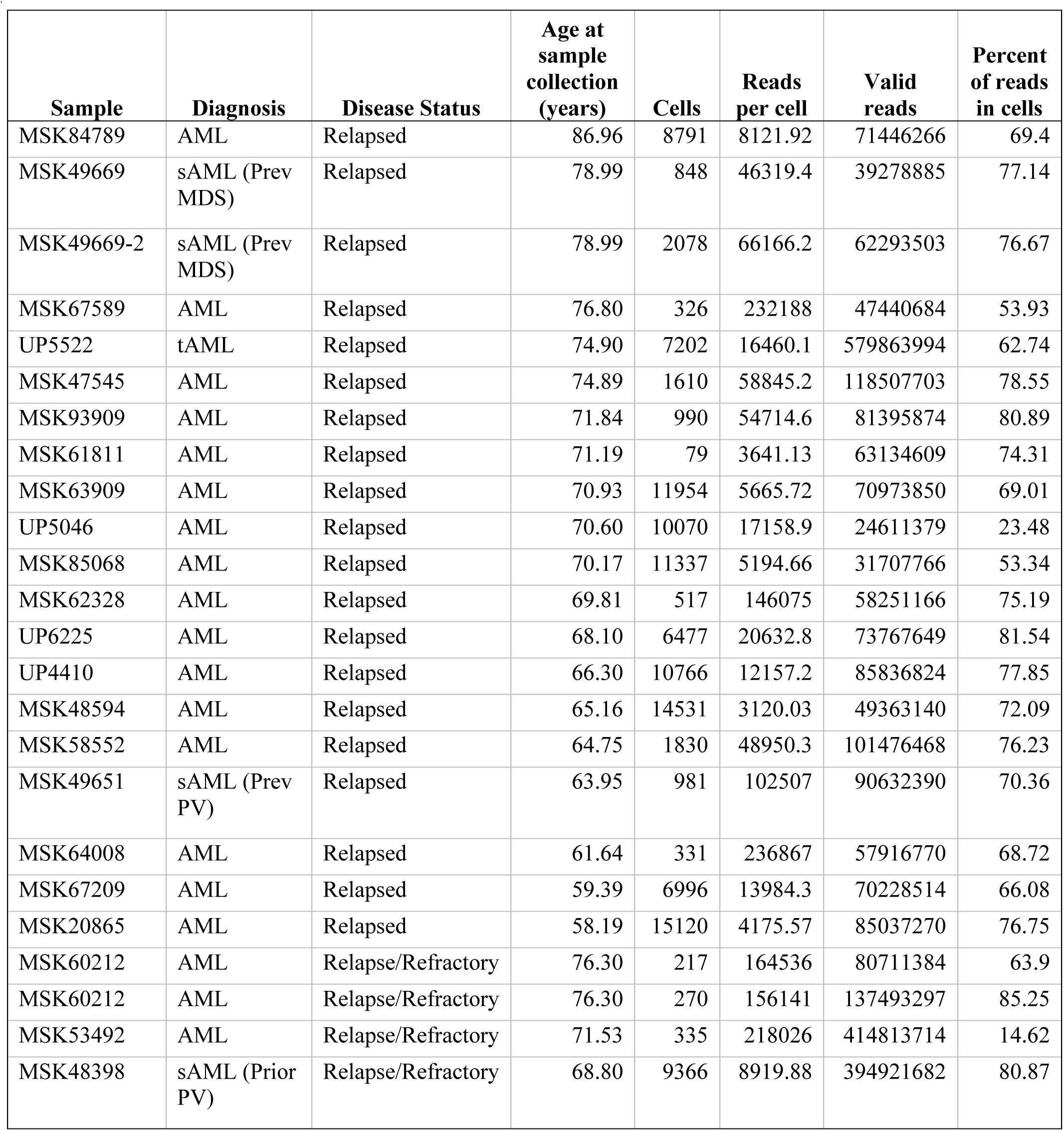
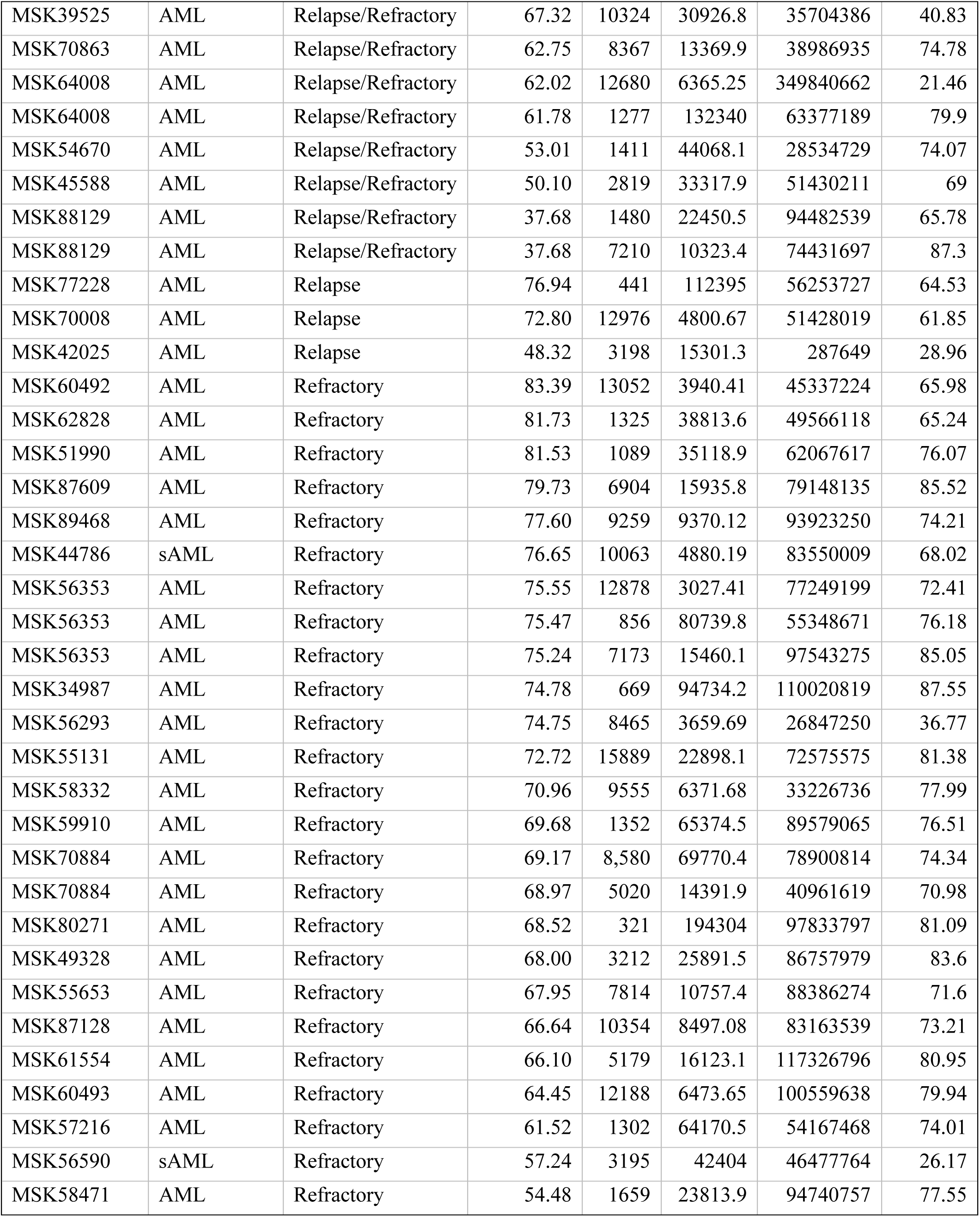
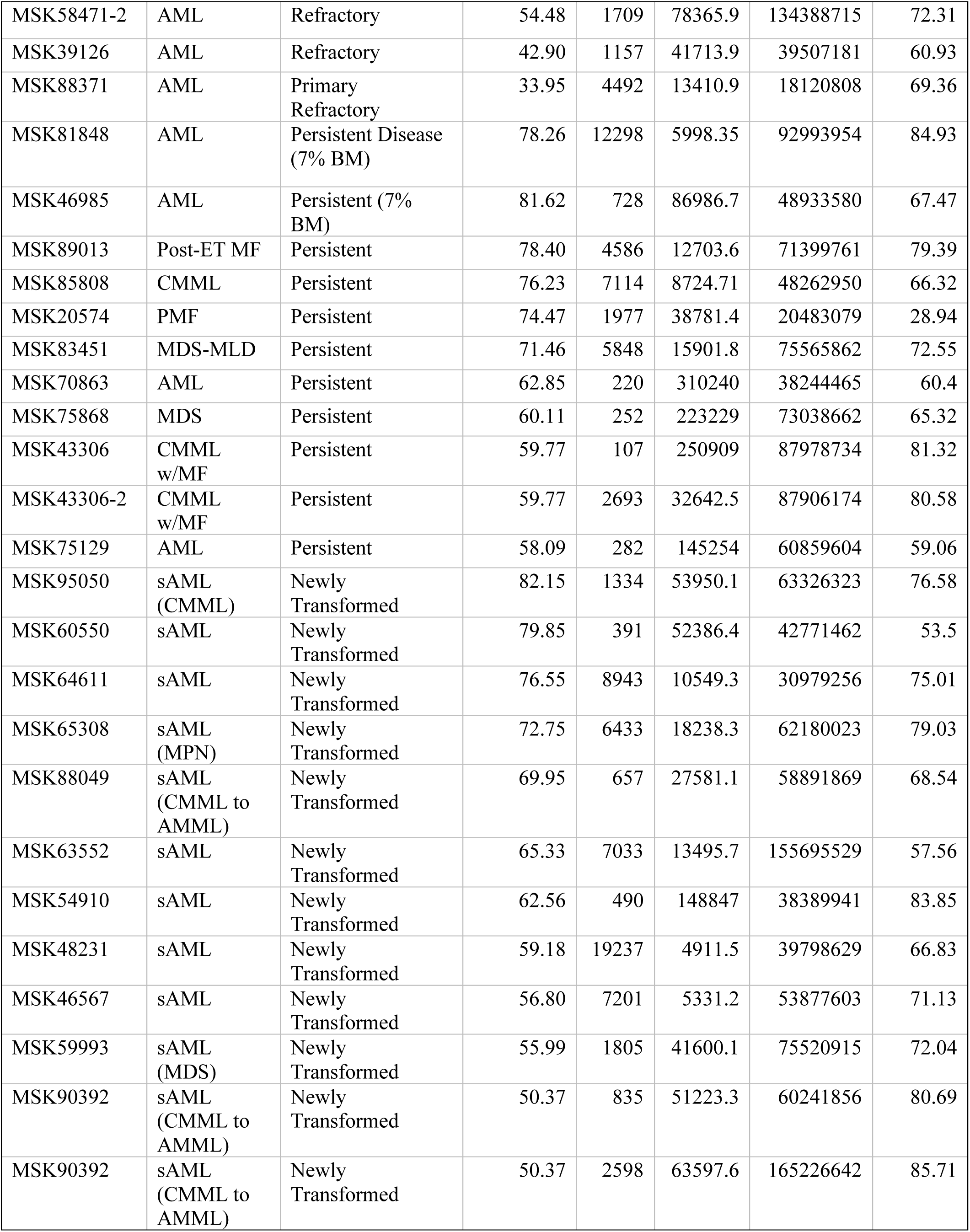
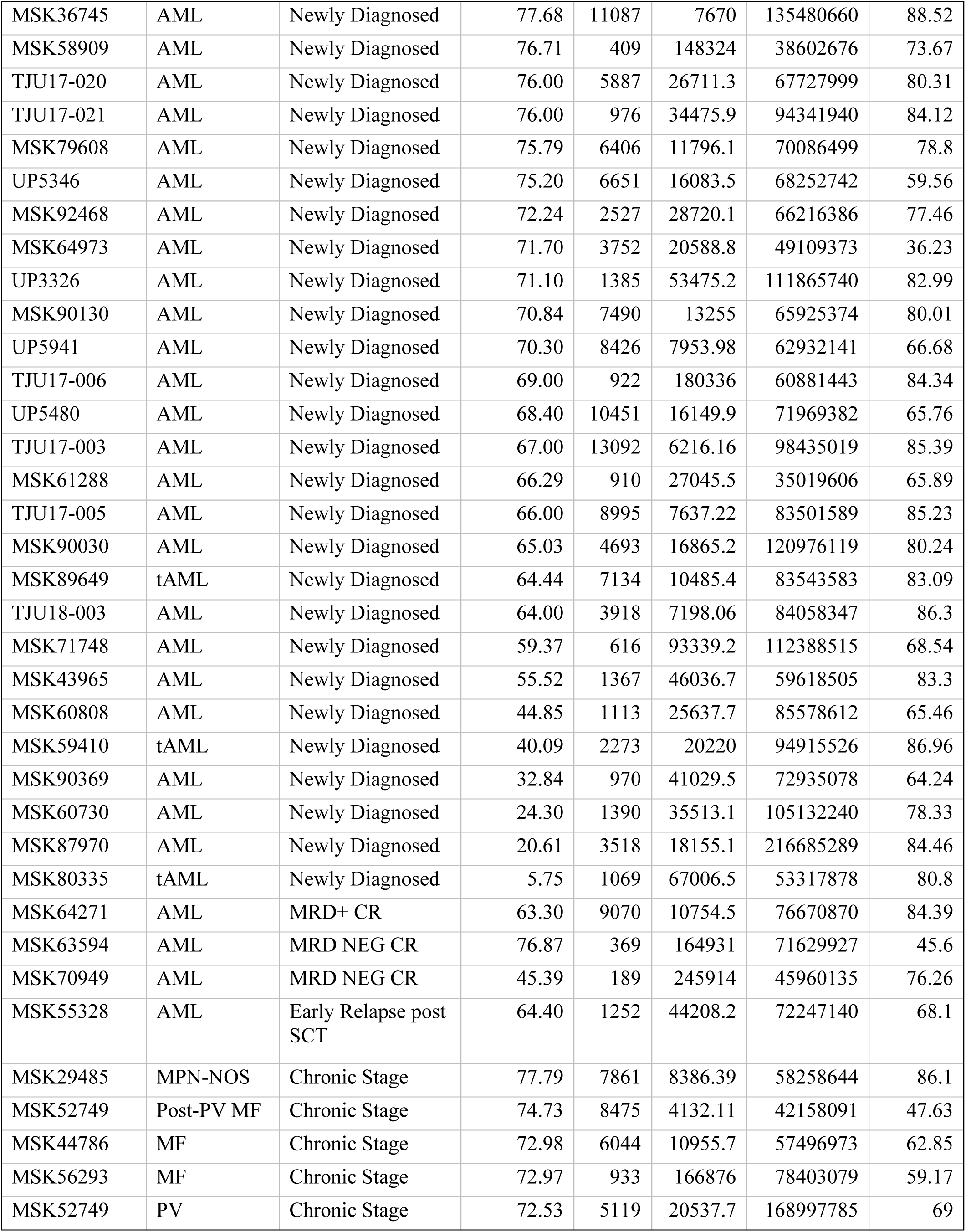
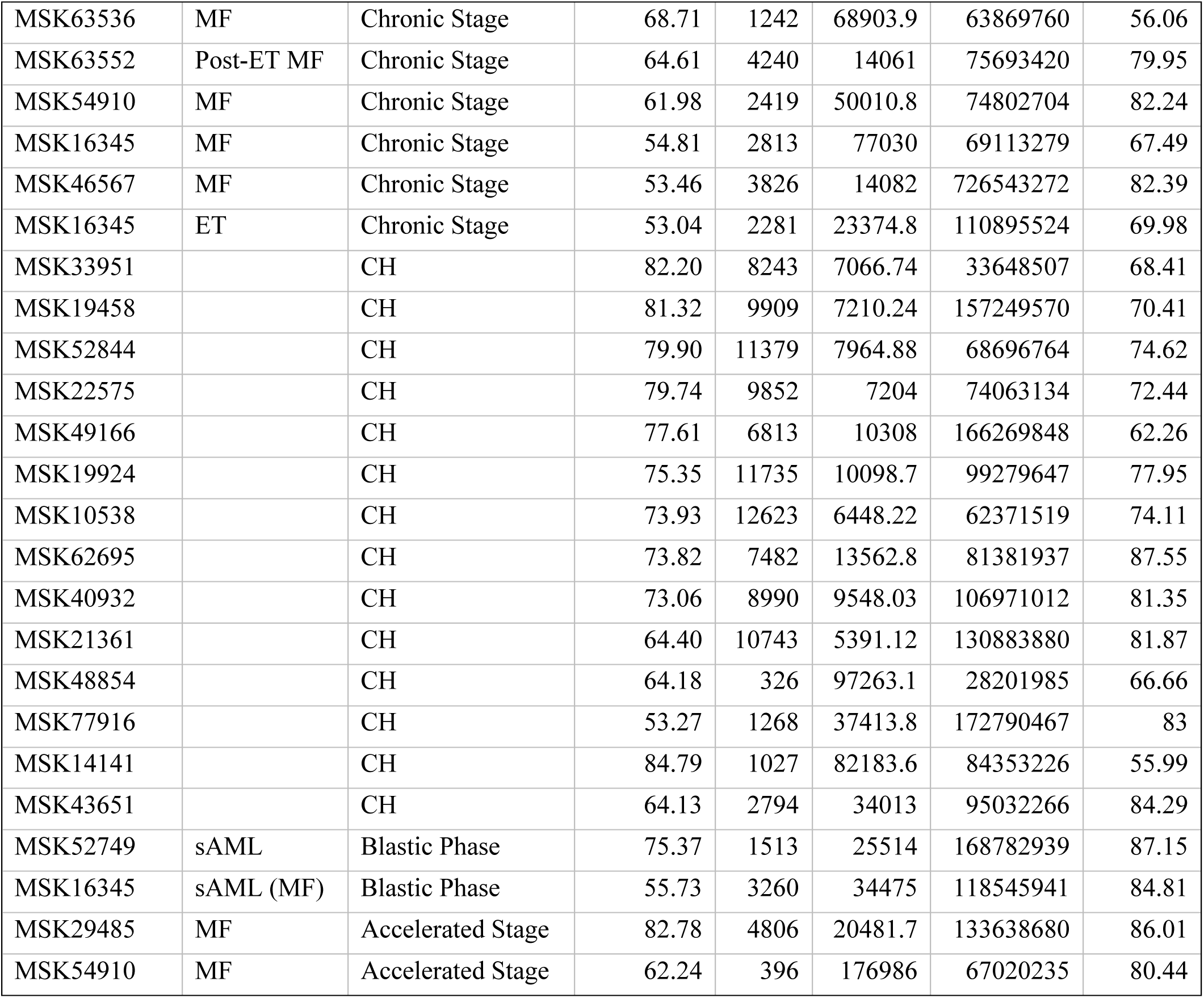
Sample characteristics from clinical cohort. Diagnosis, disease status, and age are denoted at time of sample collection. Cells, reads per cell, valid reads, and percent of reads in cells are extracted from Tapestri’s analytical pipeline on BlueBee.

**Extended Figure 1.**
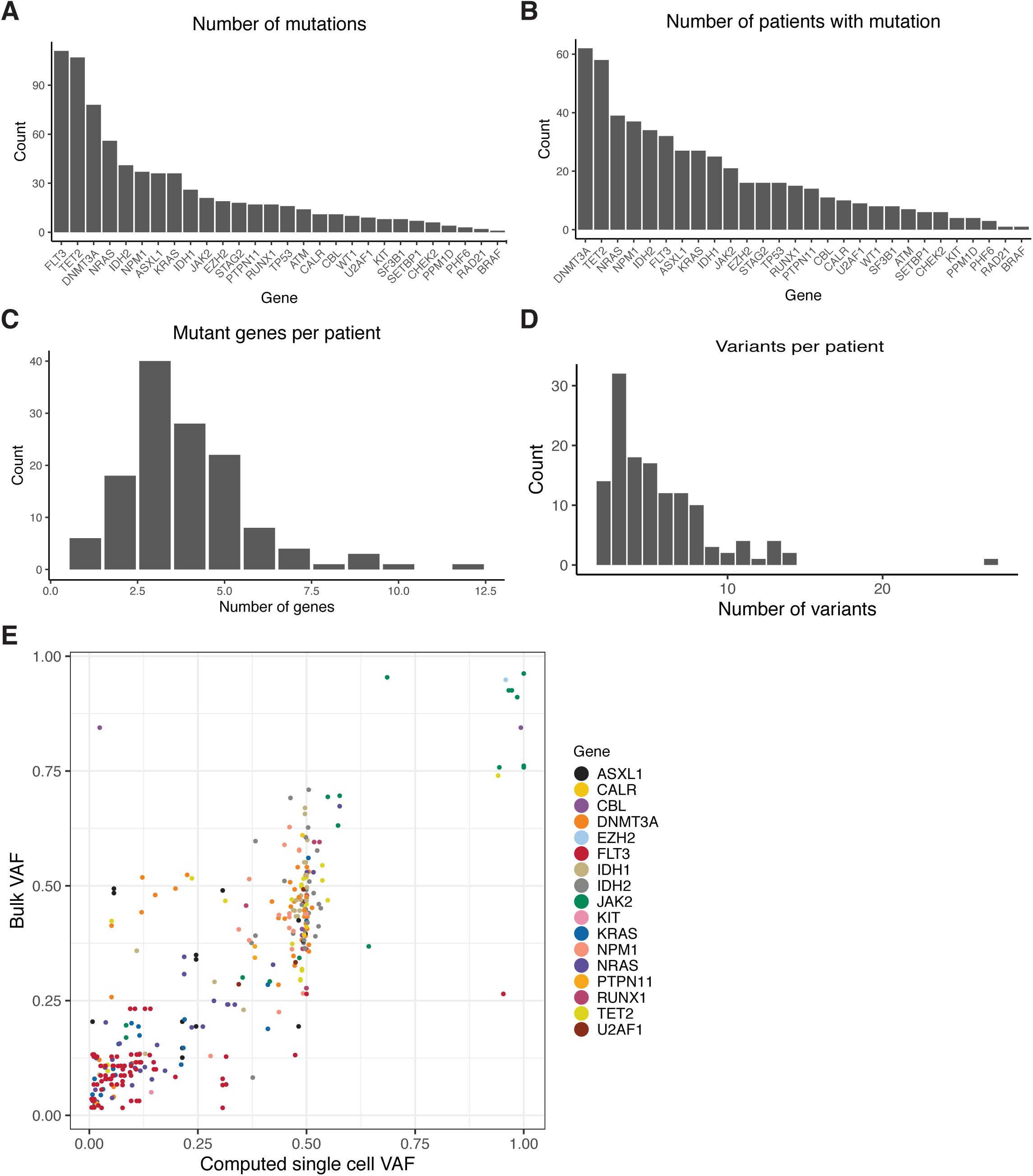
**A)** Number of individual mutations identified for each gene covered on our custom amplicon panel by single cell DNA sequencing. Genes are ranked by the number of identified protein coding mutations from highest to lowest. Genes with zero identified mutations are not listed. **B)** Number of patients with protein coding mutations in a given gene. Genes are ranked by decreasing number of patients identified with mutations. **C)** Number of patients with a given number of identified mutant genes via single cell sequencing. **D)** Number of patients with a given number of identified protein altering variants via single cell sequencing. **E)** Correlation of bulk sequencing data VAF versus single cell data VAF from MSKCC samples. Statistical significance was calculated by Pearson correlation coefficient.

**Extended Figure 2.**
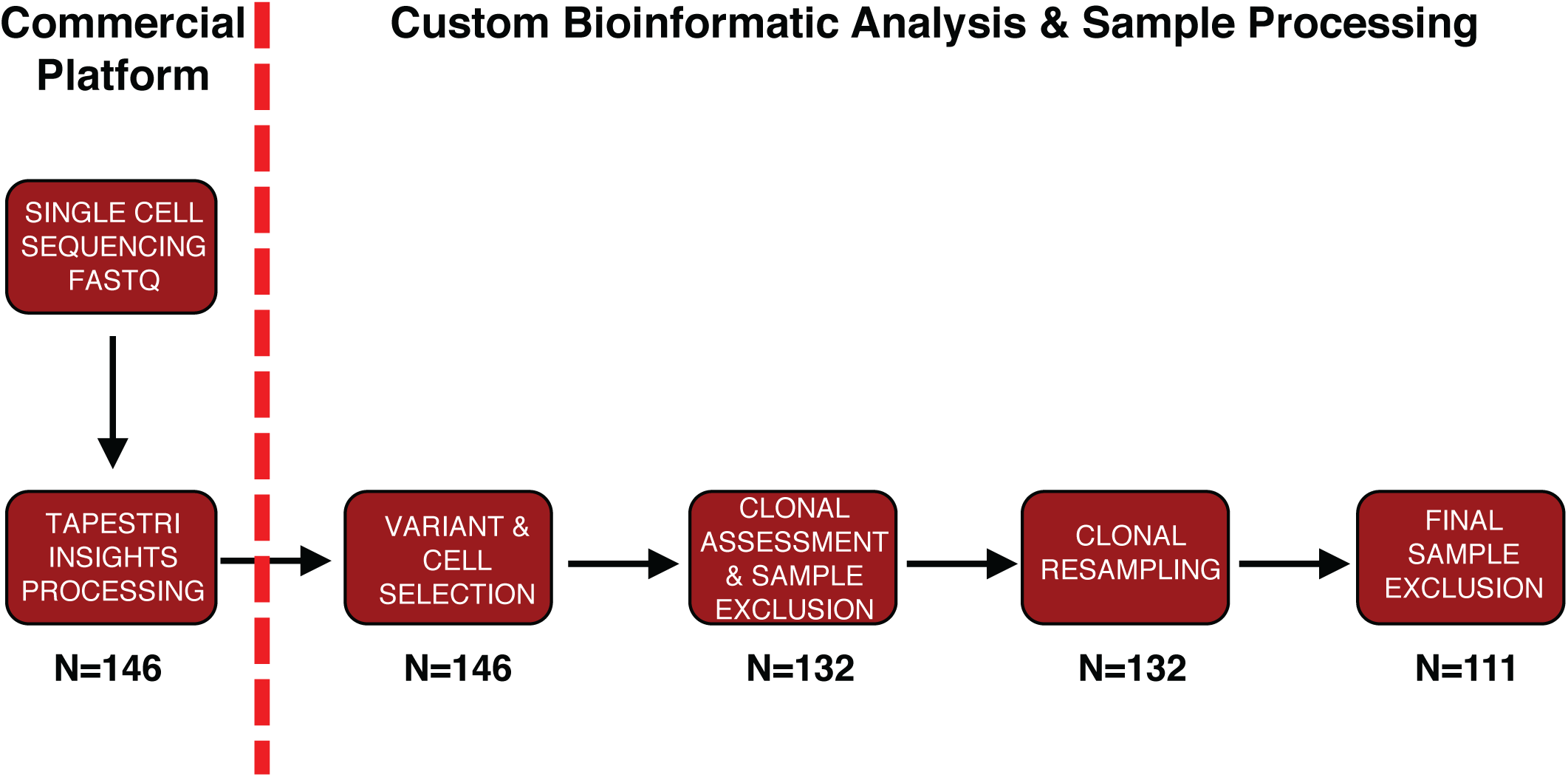
Single cell DNA sequencing data processing and analysis workflow. FASTQ sequencing files for each sample were uploaded and processed through Mission Bio Tapestri Insights platform for variant calling and cell finding (Commercial Platform). Included samples for further analysis harbored ≥1 variant which leads to a protein sequence change (non-synonymous/insertion/deletion) and included 50 cells with definitive genotyping for all protein coding variants within the sample (n=146). This data was used for analysis in Figure 1. Clones present in each sample were identified and samples removed if they contained less than 2 clones for clonal analysis studies. Samples were subjected to random resampling of cells using a bootstrapping approach to identify the stability of identified clones (n=132). Following bootstrapping, clones with lower confidence intervals <1 were removed as were variants identified only within those clones. Samples which harbored only 1 variant or presented with <2 clones after bootstrapping analysis were removed (n=111). The number of samples at each step of processing is shown below the different steps of the workflow.

**Extended Figure 3.**
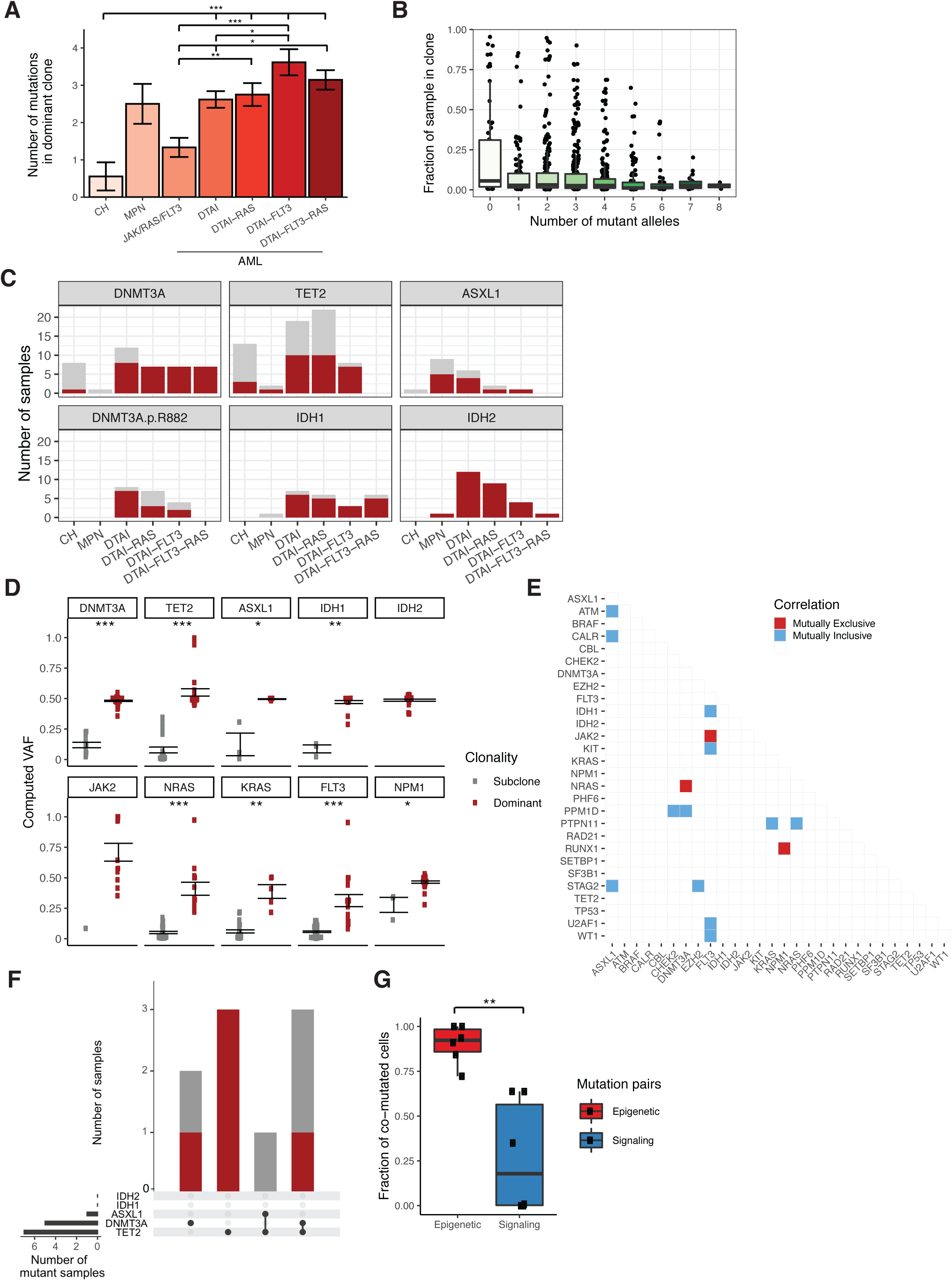
**A)** Number of mutations in the most dominant clone identified in each sample stratified by cohort. Mean value for each cohort shown by height of bar with standard error of measurement (SEM) depicted with error bars. A t-test with false discovery rate (FDR) correction was used to determine statistical significance pairwise between all groups. For clarity, only significant p-values referenced in text are shown. * P < 0.1; ** P < 0.01; ***P < 0.001. **B)** Correlation between clone size and the number of mutant alleles in the clone. Every stable mutant clone identified in clinical cohort is depicted by black circle. Mean values for each mutant allele number group shown by black bar within box with standard deviation depicted by the height of the box for each cohort. **C)** Prevalence of dominant mutations in DTAI genes across cohorts. Number of samples in each cohort with corresponding mutation. Color of bar plot annotates if mutation occurs in the dominant clone (red) or subclone (grey). Absence of bar denotes no samples were identified with mutation in a given cohort. **D)** Correlation of mutant VAF with presence or absence of mutation in the dominant clone for select genes. Computed VAF for each sample with corresponding mutation. Samples are categorized by presence of mutation in dominant clone (red) or subclone (grey). Standard error of measurement depicted with error bars. A t-test with false discovery rate (FDR) correction was used to determine statistical significance pairwise between all groups. * P < 0.1; ** P < 0.01; ***P < 0.001. Absence of p value for *IDH2* and *JAK2* due to lack of samples with subclonal mutations. **E)** Pairwise interaction matrix of mutually exclusive (red square) and inclusive (blue square) on a per sample basis. Pairwise interactions with no color did not garner a significant p-value. **F)** Lack of co-occurring DTAI mutations in CH samples. Upset plot of co-occurring DTAI mutations in CH samples with more than 1 DTAI variant. Bar graph (top panel) depicts the number of samples with each mutant gene(s) and color of bar annotating whether mutation(s) occur in the dominant clone (red) or subclones (grey). Black circles and connecting line in bottom panel demark the combination of mutations in each corresponding bar plot. **G**) Divergent frequency of co-mutated cells for epigenetic modifier genes (red) and signaling genes (blue). Individual samples shown with black square with mean denoted by black line within colored box. A Student’s t-test was used to determine statistical significance * P < 0.1; ** P < 0.01; ***P < 0.001.

**Extended Figure 4.**
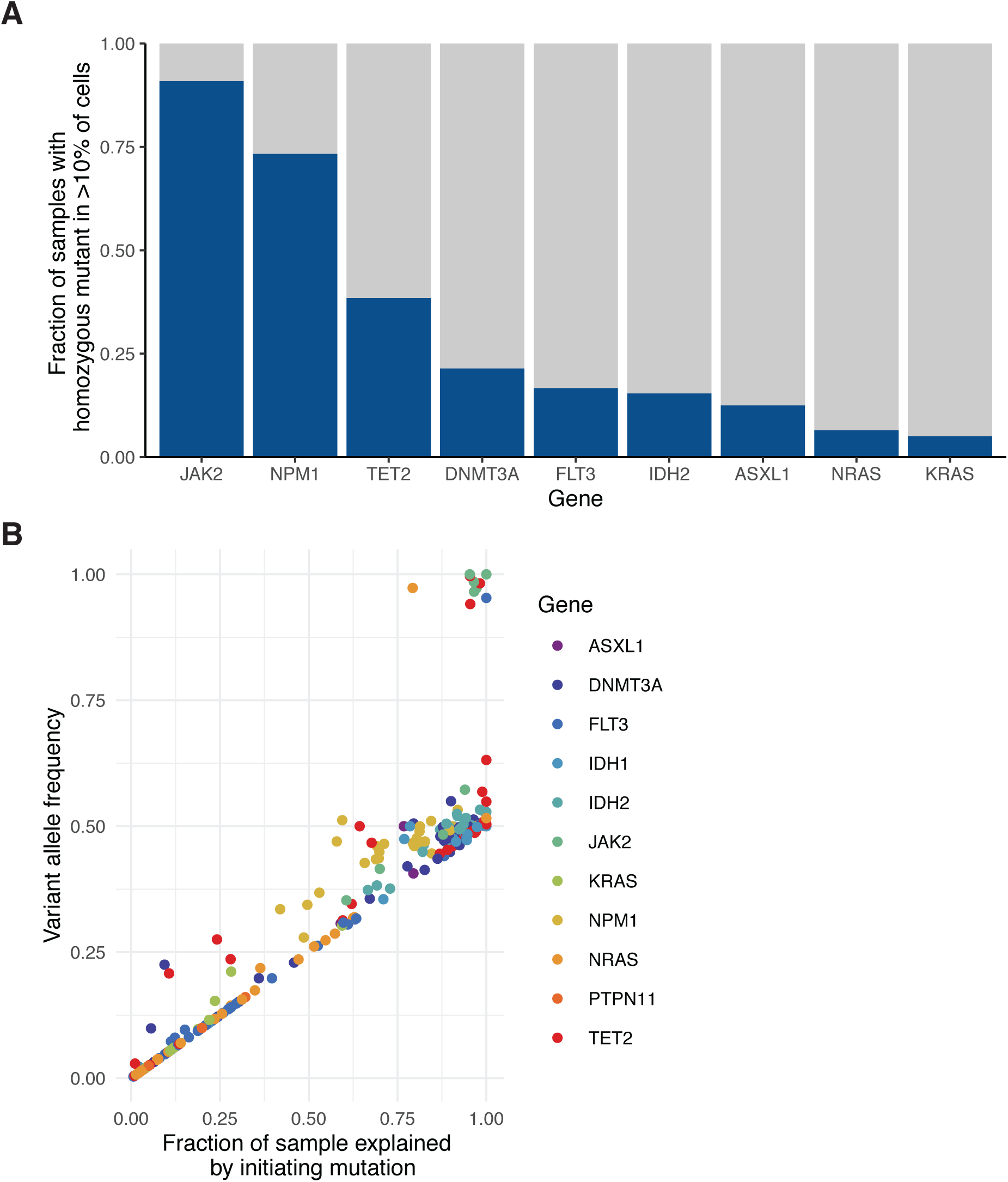
**A)** Frequency of homozygous mutations observed in all samples. Fraction of samples with homozygous mutations in a given gene (>10% of cells) based on single cell DNA sequencing data denoted in dark blue. Select genes shown and ranked by decreasing fraction of samples from left to right. **B)** Correlation of VAF computed by scDNA sequencing to fraction of a mutant sample explained by the genetic trajectory starting with an initiating mutation in a given gene. Genes used as the initiating mutation for a given sample are denoted by colored squares (colors described in figure). Statistical significance calculated by Spearman’s rank correlation coefficient test (ρ = 0.93; p ≤ 2.2 × 10^−16^).

**Extended Figure 5.**
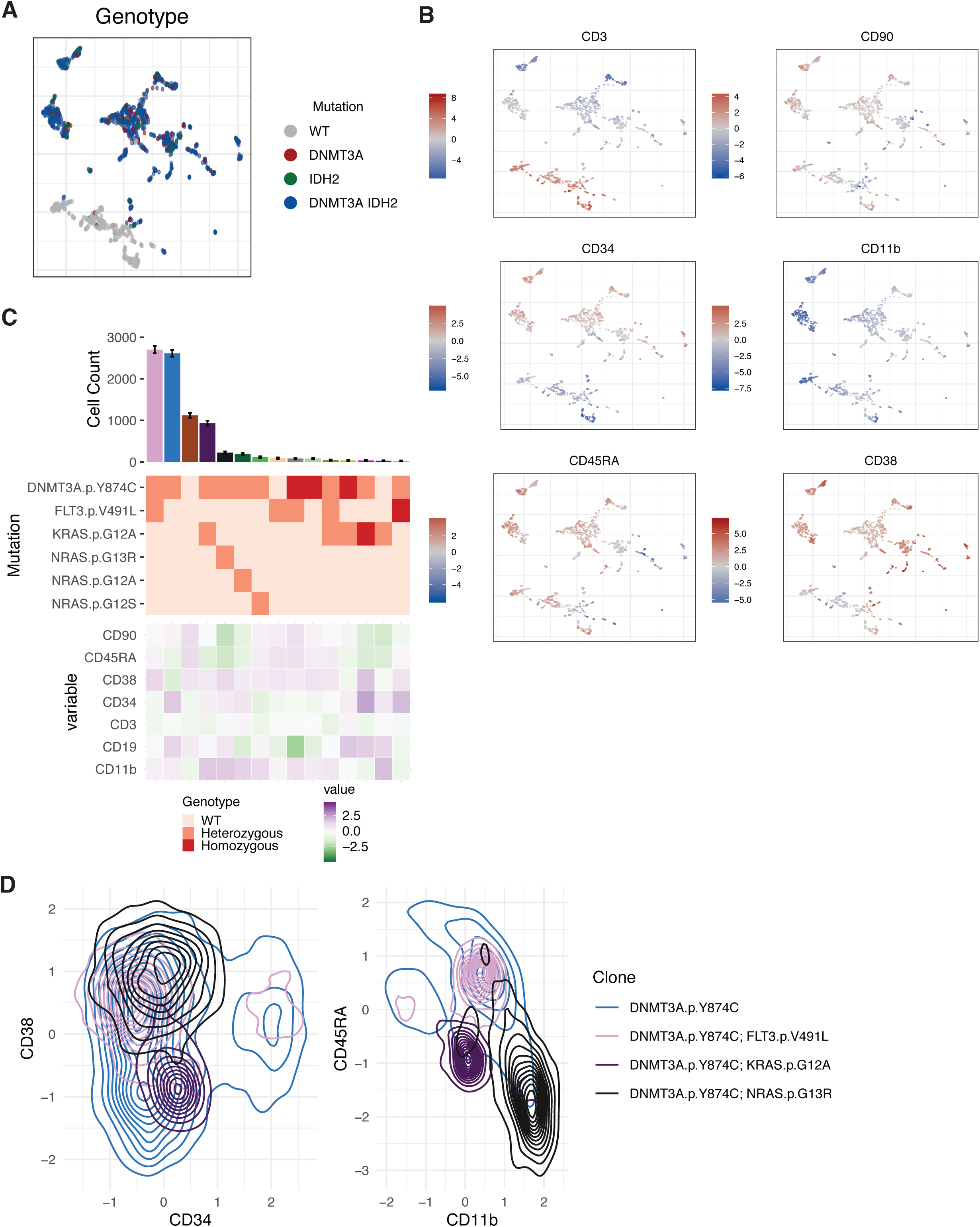
**A)** UMAP plot of MSK49669 with cells clustered by immunophenotype and genotype (WT= grey; *DNMT3A* = red; *IDH2* = green; *DNMT3A/IDH2* double mutant = blue) overlaid onto each cell. **B)** UMAP from 5A with heatmap of protein expression (high expression = red; low expression = blue) for each of the 6 antibody targets (CD3, CD11b, CD34, CD38, CD45RA, CD90) overlaid onto each cell. Relative protein expression is normalized across individual sample by centered log transformation (CLR). **C**) Clonal architecture analysis using single cell DNA + Protein sequencing of representative sample MSK56353. Clonotype plot depicts the number of cells identified with a given genotype and ranked by decreasing frequency (top panel). Cell counts for each clone is depicted with confidence intervals derived from random resampling analysis. Heatmap (middle panel) shows the genotype of each identified protein coding mutation in the given clone with zygosity (wildtype = light pink, heterozygous = orange, homozygous = red). Heatmap of the relative protein expression for each cell surface protein (n=7) in each identified clone (purple = high expression; green = low expression). **D)** Divergences in cell surface protein expression of CD34, CD38, CD11b, and CD45RA determined by presence of signaling effector mutation. Flow density plots of cells from MSK56363 of *DNMT3A* mutant cells (blue = single-mutant) with co-occurring *FLT3* (pink), *KRAS* (purple), or *NRAS* (black) mutations. Concentration of cells with a given immunophenotype depicted by the density of lines.

**Extended Figure 6.**
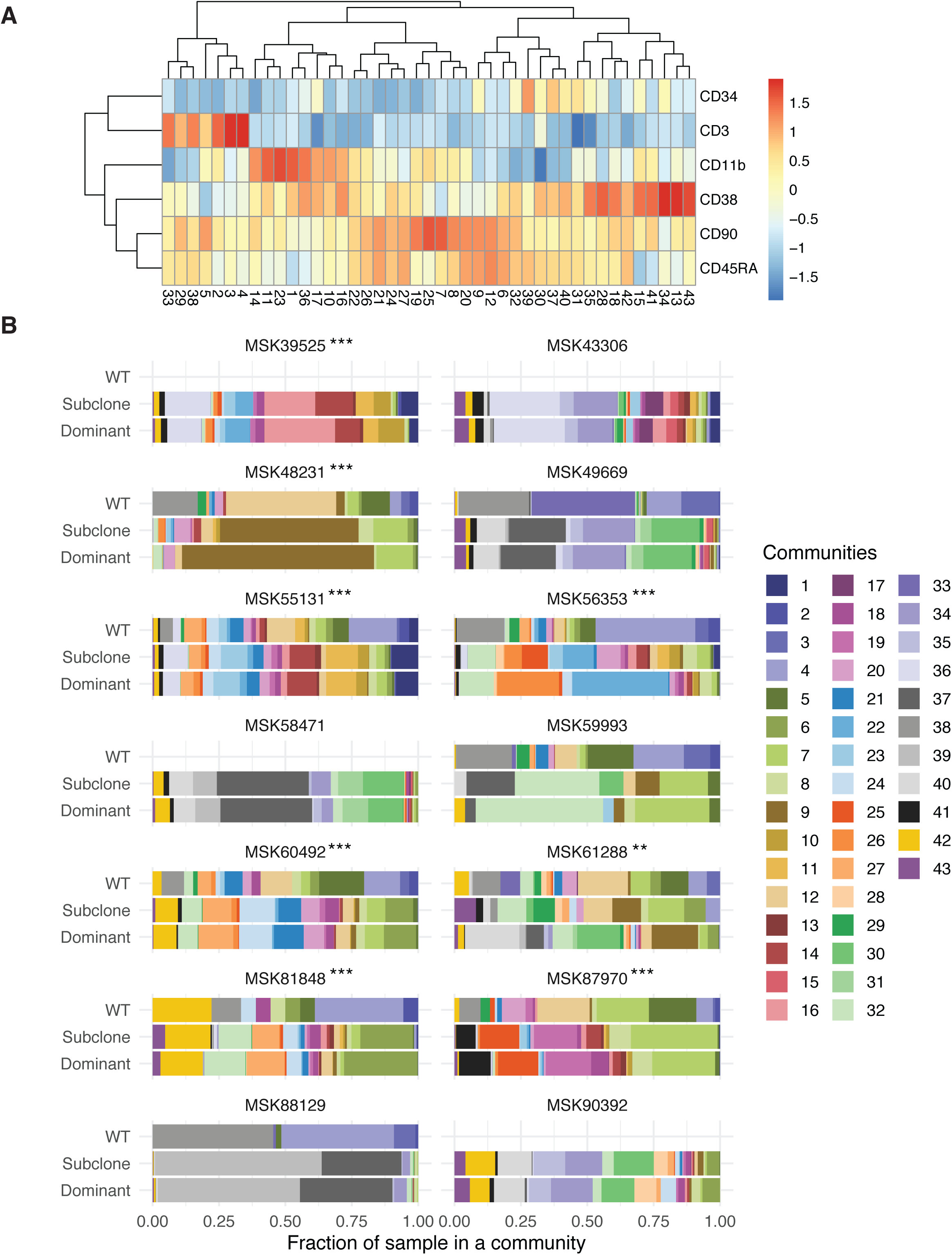
**A)** Column normalized heatmap of cell surface protein expression across 6 cell surface proteins for each community identified in phenoGraph analysis on UMAP from Figure 4H. Expression is depicted by color with blue being low expression and red annotating high expression. **B)** Community representation changes across all samples (n=14) in WT, the dominant clone, and all subclones. The fraction of each sample within each community is shown with communities depicted by corresponding color. Samples without communities shown for WT cells were found to not have any WT cells present in analysis. Changes in immunophenotype due to community representation changes for samples MSK61288 (*p* ≤ 9.95 × 10^−3^) and MSK87970 (*p* ≤ 2.45 × 10^−8^) are highlighted in Figure 4I. A two proportions z-test for each sample was used to determine statistical significance between dominant clone communities and communities present in subclone * P < 0.1; ** P < 0.01; ***P < 0.001.

